# Adaptation of human cell populations to different levels of centriole amplification

**DOI:** 10.1101/2024.02.15.580424

**Authors:** Catarina Peneda, Maria Sintra, Claudia Bank, Marco António Dias Louro, Mónica Bettencourt-Dias

**Affiliations:** Instituto Gulbenkian de Ciência/Gulbenkian Institute of Molecular Medicine, Oeiras, Portugal; Institut für Ökologie und Evolution, Universität Bern, Bern, Switzerland; Swiss Institute of Bioinformatics, Lausanne, Switzerland; Instituto de Ciências Biomédicas Abel Salazar, Universidade do Porto, Porto, Portugal

## Abstract

Centrioles are the main components of cilia and centrosomes, which play a central role in cell division and signalling. Their numbers are strictly regulated. Centriole amplification, or the presence of extra centrioles, often occurs in tumours and leads to aneuploidy and altered signalling and has been associated with cancer development and malignancy. Negative selection of cells with extra centrioles prevents numerical errors from expanding in the population, resulting in an overproduction-selection balance. However, how chronic perturbation of key centriolar regulators affects centriole number dynamics is poorly described. PLK4, a key regulator of centriole biogenesis, is often overexpressed in cancer. Here, we studied the long-term dynamics of cell populations exposed to different levels of PLK4 overexpression. We measured absolute and relative fitness in the evolving populations, quantified centriole numbers over time, as well as various aspects of the immediate response to centriole amplification. Our experiments indicated negative selection against cells with extra centrioles and outcompetition of PLK4-overexpressing cells by a cell line carrying a truncated form of PLK4, that does not amplify centrioles. In populations where cells carrying the truncated form of PLK4 were absent, cells overexpressing full-length PLK4 maintained the capacity to amplify centrioles over the course of experimental evolution and, strikingly, converge to the same degree of centriole amplification regardless of the level of PLK4 overexpression. Our results support a population-level response to centrosome amplification to control centriole amplification levels. Future work is necessary to further characterise this response and the mechanisms that allow cell populations to maintain centriole amplification.

## INTRODUCTION

The centrosome, found in most animal cells, plays a key role in organising microtubules, cell polarity, signalling, and proliferation. The canonical centrosome consists of two centrioles and a surrounding pericentriolar matrix that nucleates microtubules. During the cell cycle, each centriole duplicates once, so that in mitosis, each pair of centrioles migrates to opposite poles of the cell and directs the organisation of a bipolar mitotic spindle^1–5^. Abnormal centriole numbers can induce several changes in cell physiology. During mitosis they can cause mitotic defects that lead to chromosome segregation errors and aneuploidy and can affect cell cycle progression and viability^6–10^. In interphase, they are associated with altered secretion and signalling which can promote invasiveness^11–13^. Centriole amplification is known to be widespread in human tumours and has been associated with poor prognosis in several types of cancer, including breast cancer^14–17^. Moreover, centriole amplification is already present in pre-malignant conditions suggesting a possible role in early cancer development^18^. Furthermore, centriole amplification was shown to promote tumour initiation in mouse models^19,20^. Understanding the causes of centriole amplification and how cells respond to this perturbation is therefore vital for exploring its role in cancer development.

Centriole number is highly controlled in most cells. Even cancer-derived cell lines, which show different baseline levels of centriole amplification^21^, usually display a characteristic distribution of centriole numbers per cell. When these levels are transiently altered, centriole numbers usually revert to their original distribution^10,22–25^. This suggests that the processes that lead to an increase or decrease in the number of centrioles eventually balance each other out, producing a stable equilibrium^26^.

Centriole amplification can occur through various mechanisms, such as changes in the concentration or activity of centriolar components^27–31^, or due to failure in cytokinesis^32^. Cells with too many centrioles might divide abnormally and/or undergo cell cycle arrest, leading to reduced proliferation or even cell death. Thus, centriole numbers appear to be maintained at an equilibrium by a balance of centriole overproduction and negative selection acting on cells with extra centrioles^26,33^. However, most studies are limited to investigating short-term effects and transient centriole number alterations. The long-term dynamics of centriole numbers, especially in cancer where centriole-related proteins are often overexpressed^34,35^, remain poorly understood. Thus, understanding these dynamics in scenarios of chronic perturbation is crucial for a comprehensive view of centrosome regulation.

PLK4, an essential centriole biogenesis regulator, plays a significant role in centriole number dynamics. Several studies showed that overexpressing PLK4 leads to centriole amplification, whereas reducing its expression can prevent centriole duplication^27,28^. In cultured cells, short-term chronic PLK4 overexpression leads to proliferation defects that limit the proliferation of cells bearing extra centrioles^6,36^. In mice, it was proposed that mild PLK4 overexpression is associated with persistent centriole amplification and cancer development^19,20^. In contrast, it was suggested that cells with high levels of expression are negatively selected^36–38^. However, this hypothesis has not been tested. Although PLK4 overexpression is common in cancers like breast cancer and acute myeloid leukaemia^39–41^, where centriole amplification has been observed, the impact of these gene expression changes on centriole number dynamics is not known. This study aims to explore how different PLK4 overexpression levels affect centriole number dynamics and cell population fitness over time.

We used MCF10A cells carrying a doxycycline-inducible system to trigger PLK4 overexpression^11^ at different levels. Over two months we monitored centriole numbers, conducted competition assays for fitness and proliferation, and tracked PLK4 levels.

## RESULTS

### An experimental framework for addressing the evolution of cell populations experiencing different levels of centriole amplification

In this study, we established an experimental framework for addressing the long-term overproduction-selection dynamics of populations with centriole amplification. We made use of overexpression of PLK4, as the trigger for centriole amplification. We chose to use MCF10A-PLK4, a cell line derived from non-transformed breast tissue^11^, where PLK4 overexpression is induced by adding doxycycline (Dox) to the culture medium. This system is widely used to study the implications of altered centrosome number in cell physiology and breast cancer^11,13,21,42,43^, widening the implications of our study.

We first sought to obtain cell populations with different initial degrees of centriole amplification. Tetracycline/doxycycline-inducible systems are known to display a sigmoidal dose response - i.e. the expression of target gene(s) rises sharply for intermediate levels of induction lead and plateaus when higher doses are applied^44^. This behaviour led us to search for optimal intermediate doses, which would lead to different levels of centriole amplification. Previously, we had observed that 24h treatment with doxycycline concentrations between 0.1 μg/mL and the typically used 2 μg/mL, yielded similar levels of centriole amplification (data not shown). Thus, we investigated the effect of treating MCF10A-PLK4 populations with concentrations of Dox ranging between 0 and 0.1 μg/mL for 24h (Fig. 1A, B). Since centriole numbers change along the cell cycle, we analysed mitotic cells, which are expected to have four centrioles, allowing us to estimate the degree of centriole number abnormalities more accurately. We observed a significant and dose-dependent increase in the relative frequency of mitotic cells with extra centrioles in Dox-treated populations compared to the control without Dox. Treatment with 0.001 μg/mL of Dox resulted in approximately 29% of cells with extra centrioles. Between 0.005 and 0.1 μg/mL of Dox, the relative frequency of mitotic cells with extra centrioles plateaued at around 60-70% (Fig. 1B). Given that we observed significant differences in the level of centriole amplification in populations treated with Dox at 0, 0.001, and 0.1 *μ*g/mL, we selected these concentrations for further investigation. We will refer to them hereafter as Dox0, Dox0.001 and Dox0.1.

**Figure 1.**
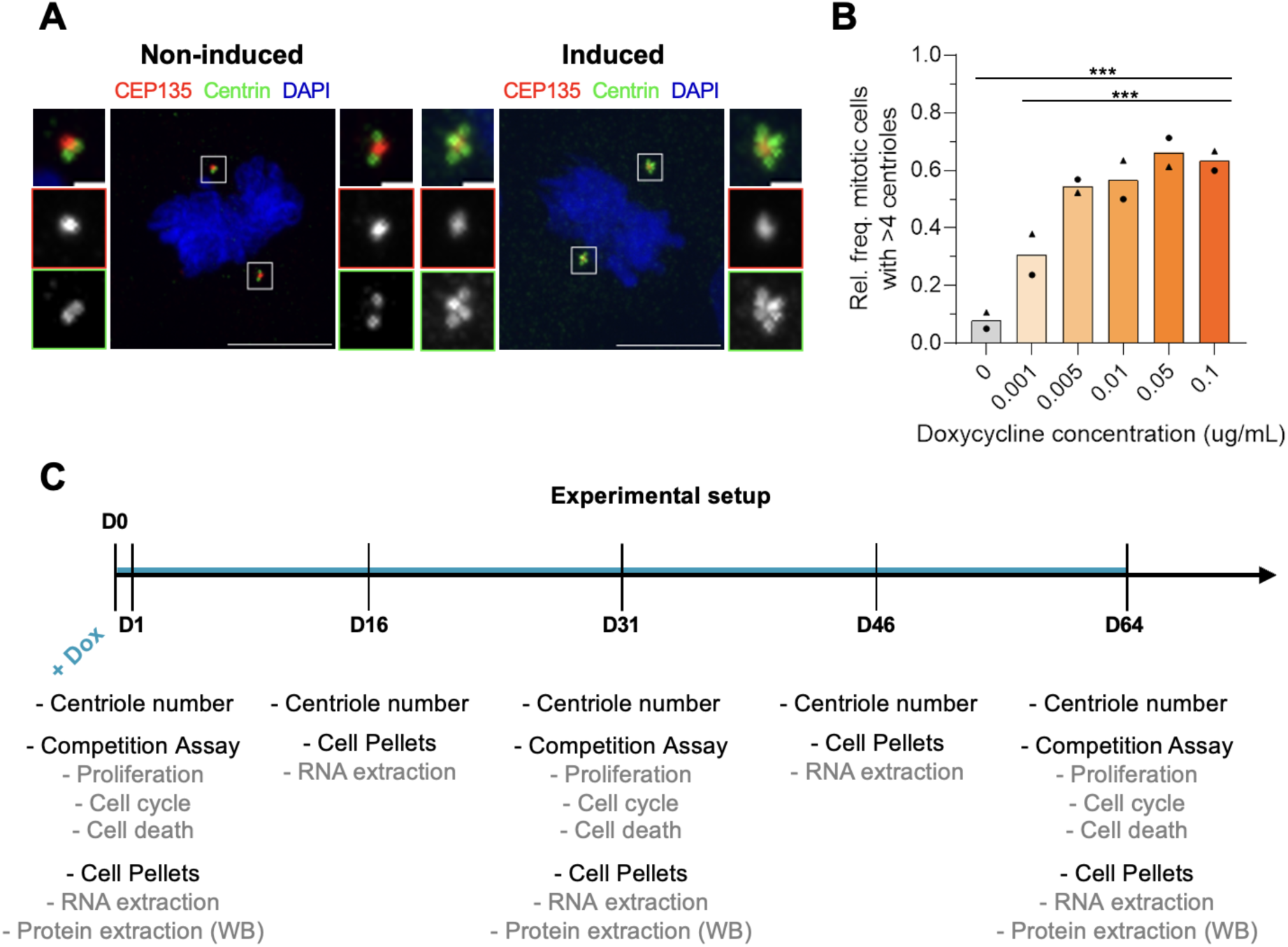
Experimental framework for addressing the evolution of cell populations experiencing different levels of centriole amplification. **A** - Example of mitotic cells with wild-type (left) and abnormally high (right) centriole numbers. Cells were stained for CEP135 (magenta), Centrin (green), and DNA (blue). Scale bar is 10 nm. B - Relative frequency of mitotic cells with more than four centrioles following 24h treatment with the indicated doses of doxycycline. Symbols represent different replicates. 62-100 mitotic cells were analysed in each of the two independent experiments. **C**- Experimental evolution setup and conditions. MCF10A-TetR, MCF10A-PLK4^1−608^ and MCF10A-PLK4 populations were grown for two months in media containing doxycycline at 0, 0.001, or 0.1 μg/mL and sampled for centriole number counting, competition assays, and pellets for RNA extraction and western blots, at the indicated time points.

To assess the consequences of long-term chronic PLK4 overexpression and centriole amplification, we designed an experimental evolution setup (Fig. 1C). Experimental evolution aims at characterising how evolutionary forces, such as natural selection or genetic drift, shape populations because of the established test conditions^45^. We were interested in testing if and how cell populations subject to chronic PLK4 overexpression adapt to centriole amplification and how this might occur. MCF10A-Plk4 is a polyclonal cell line, thus containing genetic and phenotypic variation that may allow cells to cope with PLK4 overexpression or centriole amplification. Second, we evolved cells for two months (approximately 90 generations according to published doubling times for regular MCF10A in control conditions), which is sufficient for selection to act while preventing excessive accumulation of mutations. Third, we passaged cells every three days at a high seeding density (2.2×10^6^ cells) and allowed them to reach a large population size (∼2×10^7^ cells in control conditions). This setup provided us control over the population growth dynamics and enabled us to minimise genetic drift.

Given that both Dox alone^46^ and PLK4 catalytic activity^47^ may induce physiological effects beyond centrosome amplification we added different controls to our experiment. We controlled for other consequences of expressing PLK4 and adding Dox using two different cell lines: 1) MCF10A-PLK4^1−608^, which carries a Dox-inducible truncated form of PLK4, which includes the kinase domain but not its C-terminus necessary to locate it to the centriole. Thus, its overexpression does not result in centriole overproduction^11,48^ and 2) MCF10A-TetR, the parental cell line of MCF10A-PLK4 and MCF10A-PLK4^1−608^, which carries the Dox-inducible system but not a target transgene. Thus, our setup included three different cell lines in three different cell culture media (containing 0, 0.001, or 0.1 μg/mL of Dox), which we evolved for two months (Fig. 1C).

This setup allowed us to assess a variety of cellular parameters over time. We sampled MCF10A-TetR, MCF10A-PLK4^1−608^, and MCF10A-PLK4 populations grown with 0, 0.001, or 0.1 *μ*g/mL of Dox at days 1, 3, 16, 31, 46, and 64 for centriole number counting and RNA extraction. In addition, we performed competition assays in the beginning (starting at day 1), middle (starting at day 31) and at the end of the experiment (starting at day 64). This enabled us to estimate population fitness over time, as well as to assess cell proliferation, cell death, and cell cycle profiles using flow cytometry. Finally, we monitored population growth using automated cell counting during passaging.

### Centriole number dynamics depend on the initial centriole amplification levels in MCF10A-PLK4

We first asked how centriole numbers change in cell populations chronically treated with different doses of doxycycline. As expected, the relative frequency of cells with extra centrioles showed no significant differences over time in the negative controls MCF10A-TetR and MCF10A-Plk4^1−608^, regardless of Dox (Fig. 2A-B, Fig. S1), and in non-induced MCF10A-PLK4 (Fig. 2C). In contrast, the relative frequency of mitotic cells with centriole amplification in Dox0.001- and Dox0.1-treated MCF10A-PLK4 reached an average of 28±4% and 91.5±5.5% after 24h, respectively, compared with 7±2% for the non-induced population, and steadily decreased in subsequent days (Fig. 2C). The number of centrioles per mitotic cell was highest, on average, and most variable at day 1 (mean centriole number: Dox 0 - 4.18±0.02, Dox0.001 - 5.24±0.26, Dox0.1 - 12,84±2.14; Fig. 2D-F). Over time, both the mean and variance of centriole numbers per cell reverted to basal levels for both Dox concentrations (Fig. 2D-E). Finally, centriole amplification decreased more rapidly at higher Dox concentrations (Dox0 - 0.00173, *p*-value=0.97653, Dox0.001 - −0.23567, *p-*value=0.00128, Dox0.1 - −0.864769, *p*-value<2e^-16^). This suggests that different PLK4 overexpression levels/incidence of cells with extra centrioles can elicit different selective pressures. We concluded that both low and high chronic Plk4 overexpression ultimately result in loss of centriole amplification, but with different dynamics. These results suggest that populations under chronic induction of PLK4 overexpression adapted by inhibiting overproduction or that continuous negative selection eliminated cells with extra centrioles, leading to a reduction in population fitness. Therefore, we investigated the fitness of these populations.

**Figure 2:**
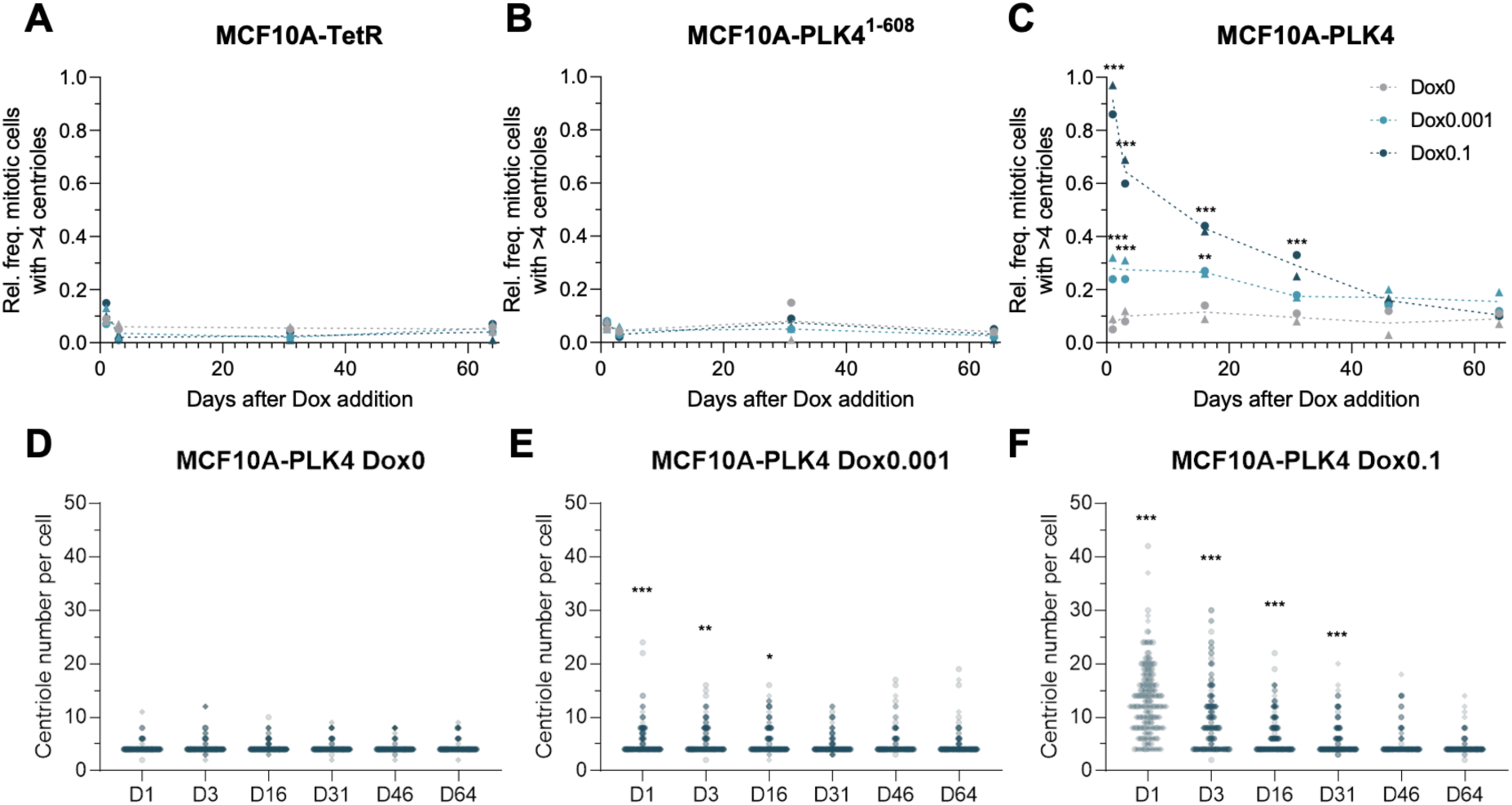
Centriole number dynamics depend on the initial centriole amplification levels in MCF10A-PLK4. **A-C-** Relative frequency of mitotic cells with more than four centrioles at the indicated time points of Dox treatment for MCF10A-TetR (A), MCF10A-PLK4^1−608^ (B), and MCF10A-PLK4 (C). Data points correspond to populations grown in the presence of doxycycline at 0 (light grey), 0.001 (light blue), or 0.1 *μ*g/mL (dark blue), dashed lines represent the average relative frequency of cells with more than four centrioles at each time point for populations grown in Dox0 (grey), Dox0.001 (light blue), or Dox0.1 (dark blue) and different shapes represent independent experiments. Statistical analysis were performed using a generalised linear model with the presence/absence of extra centrioles in a cell as a binomial response variable and taking Dox concentration and time point as independent categorical predictors, plus their interaction. Significant differences between the indicated condition and the one treated with Dox0 at the same time point are represented with ** (*p-*value < 0.01) or *** (*p-*value < 0.001). **D-F** - Centriole number distributions in MCF10A-PLK4 populations growth in Dox0 (D), Dox0.001 (E), and Dox0.1 (F). 72-100 mitotic cells were analysed for each condition in each of the two independent experiments.

### MCF10A-PLK4 populations adapted to centriole amplification in a dose-dependent manner

Estimating competitive fitness requires a setup that allows populations to be distinguished in co-culture. To accomplish this, we labelled these populations with proliferation dyes, CellTrace CFSE and CellTrace Far Red^49^.

To monitor population fitness over time, we performed competition assays in which we co-cultured MCF10A-PLK4 or the control MCF10A-PLK4^1−608^ with the parental cell line, MCF10A-TetR. Each co-cultured population was labelled with a different proliferation dye and grown for three days (Fig. 3A). This time window allows for a clear separation of both sub-populations by flow cytometry and encompasses sufficient cell divisions for detecting fitness differences (Fig. S2). As technical controls, we performed replicate co-cultures stained with the opposite combination of dyes, controlling for the effect of the dye on proliferation. Finally, we calculated the competitive index for each co-culture, a standard measure of competitive fitness^50^ (see Methods).

**Figure 3.**
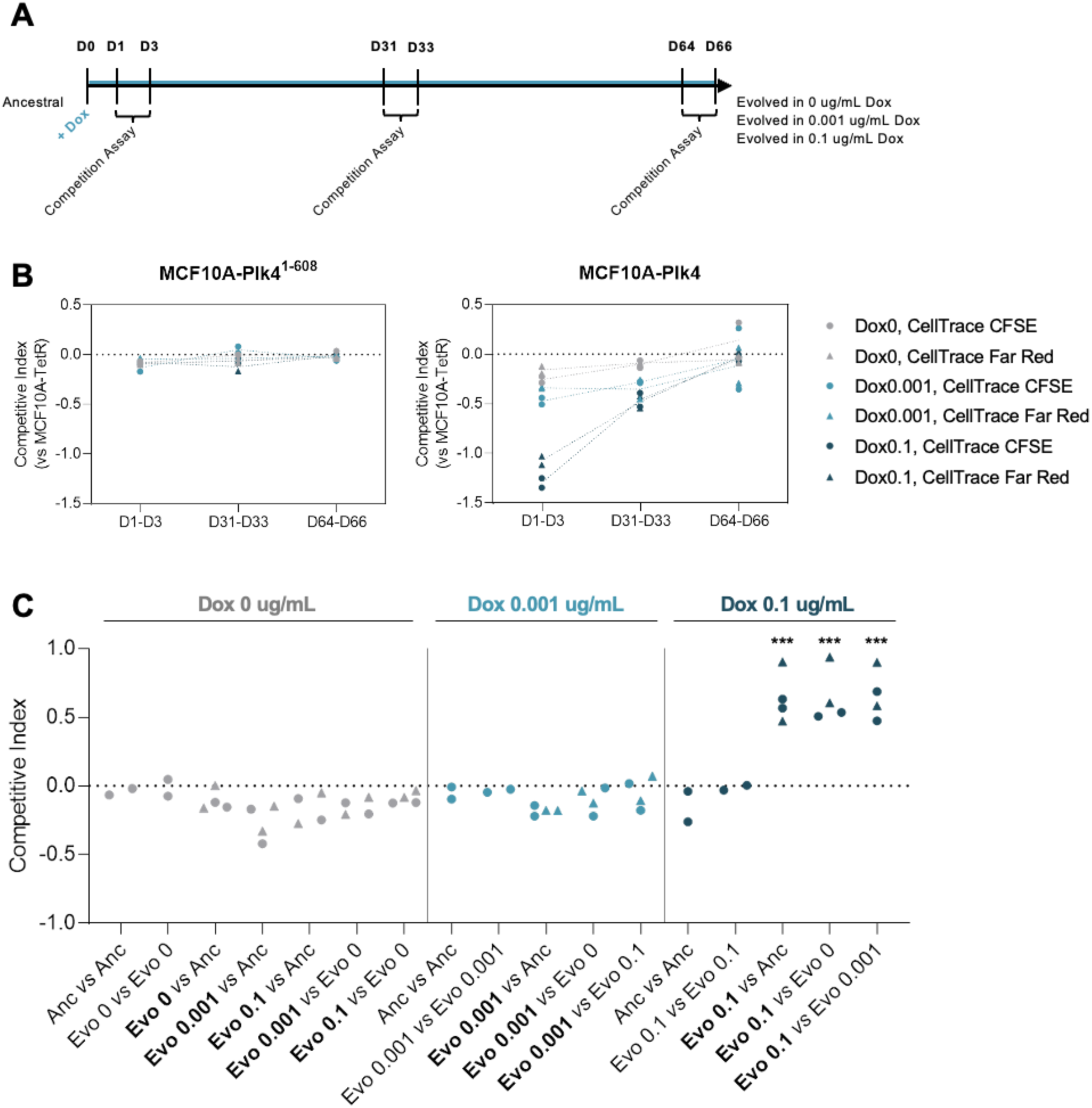
Induced MCF10A-PLK4 adapted over time to chronic PLK4 overexpression. **A -** Timeline for competition assays. Cells were labelled with different fluorescent dyes (CellTrace CFSE or Far Red) and co-cultured at the indicated time points. The competitive index was used as a proxy for competitive fitness based on the relative frequencies of each co-cultured population at the initial and final time points (see Methods). **B -** Competitive index of MCF10A-PLK4^1–608^ (B) or MCF10A-PLK4 (C) relative to MCF10A-TetR at the indicated time points. Note that for each experimental situation there are 4 data points (independent experiments and swapping of the dye). The dashed lines connect the average competitive index corresponding to each Dox concentration and dye combination in two independent experiments. See table S1 and S2 for statistical analyses. **C -** Competitive index of evolved and ancestral MCF10A-PLK4 populations competing in the indicated combinations. Cell populations were co-cultured in medium with Dox0, Dox0.001, or Dox0.1. The points in the plots correspond to the competitive index of the cell line in bold. Circles correspond to populations labelled with CellTrace CFSE and triangles to populations labelled with CellTrace Far Red. Note that when a given population was co-cultured with itself, one sub-population was labelled with CellTrace CFSE and the other with CellTrace Far Red, i.e. there is only one combination of dyes. Data were modelled using a linear model (ANOVA) taking experimental conditions as a grouping variable. Significant differences between the indicated conditions and their controls (ancestral versus ancestral and evolved versus evolved) are represented with *** (*p*-value<0.001).

Importantly, for all data points, we observed a good correlation between replicates and between different proliferation dye combinations (Fig. S2). Additionally, we observed no effect of the initial frequency of each co-cultured sub-population on competitive index estimates (Fig. S2). Together these suggest that the calculated index reflects changes in fitness and not experimental artefacts. As expected, control line MCF10A-PLK4^1−608^ showed no significant fitness differences between different Dox concentrations when co-cultured with MCF10A-TetR at any time point (Fig. 3B Left panel). This is coherent with previous reports, in which short term overexpression of truncated PLK4 did not affect population growth^11^. However, we observed that Dox treatment imparted a significant and dose-dependent fitness cost in MCF10A-PLK4 relative to MCF10A-TetR at days 1-3 (Fig. 3B right panel). At days 31-33, the fitness cost associated with both Dox concentrations was highly reduced and we found it to be undetectable at days 64-66 (Fig. 3B right panel). These results suggest that the fitness cost observed in the induced MCF10A-PLK4 populations depends on the overexpression of full-length PLK4 and/or the presence of centriole amplification.

Our results led us to ask if the evolved populations, i.e. from the end point of the experiment, had adapted to the presence of Dox. Adaptation can be defined as a relative increase in fitness of the evolved populations compared with the ancestral populations, in the conditions in which they evolved^51^. To address this, we co-cultured the evolved MCF10A-PLK4 populations with the ancestral population in media with different Dox concentrations. Moreover, to ask whether Dox concentration led to differently adapted populations, we co-cultured each pair of evolved populations in different combinations of media (Fig. 3C).

As controls, we co-cultured each evolved and ancestral populations with themselves, and observed no significant fitness differences (Fig. 3C). The population evolved in Dox0.001 did not show significant fitness differences compared with the ancestral population in any media, indicating no significant adaptation in this condition (Fig. 3C). On the other hand, the population evolved in Dox0.1 outcompeted all the others at this concentration, suggesting it had adapted (Fig.3C).

Adaptation to an environment often involves molecular changes that can have a fitness cost if cells return to the ancestral environment^52^. Interestingly, when we co-cultured each of the three evolved populations with the ancestral population, or the population evolved without Dox, in media without Dox, we observed no significant fitness differences (Fig. 3C). These results suggest a negligible fitness cost of Dox adaptation.

In summary, (1) MCF10A-Plk4 populations adapted to high (Dox0.1) but not low levels (Dox0.001) of chronically-induced PLK4 overexpression; (2) there was no cost of adaptation; (3) exposing the populations evolved in Dox0 and Dox0.001 to higher doses of Dox still bears a fitness cost. Therefore, since adaptation to centriole amplification/PLK4 overexpression was dose-dependent, we hypothesised that populations grown in different doses of Dox may have developed distinct mechanisms that allow them to cope with PLK4 overexpression/centriole amplification (e.g. regulating PLK4 expression, or even completely silencing transgene, as previously observed^37^). Notwithstanding, these putative mechanisms both resulted in the decrease of extra centrioles from the population.

Since our competition results suggest that the fitness depends on the level of centriole amplification, we asked if this could also explain the observed fitness differences between evolved populations. To address this, we quantified centriole numbers in each of the evolved populations treated with Dox0, 0.001, and 0.1 for 24h. Moreover, we tested if they could produce extra centrioles when treated with higher concentrations of Dox (2 μg/mg). As controls, we treated ancestral populations with the same range of Dox concentrations.

We observed that the ancestral MCF10A-PLK4 populations showed a proportional increase in the relative frequency of mitotic cells with extra centrioles when treated with 0.001 and 0.1 μg/mL of Dox for 24h (Fig. 4A, B), as observed above. Treatment with 2 μg/mL of Dox yielded no significant differences compared with 0.1 μg/mL. The populations that evolved in Dox0 showed an identical trend compared with the ancestral population but reached lower levels of amplification for each respective treatment. Populations evolved in Dox 0.001 did not show a response when grown in that medium or without Dox. However, they showed an increase in the relative frequency of cells with extra centrioles when exposed to 0.1 and 2 μg/mL of Dox. Concerning the population that evolved in Dox 0.1, none of the tested treatments yielded a significant increase in the relative frequency of mitotic cells with extra centrioles (Fig. 4A, B). This suggests that the mechanism of inhibition of centriole overproduction may be specific to the level of PLK4 overexpression.

**Figure 4.**
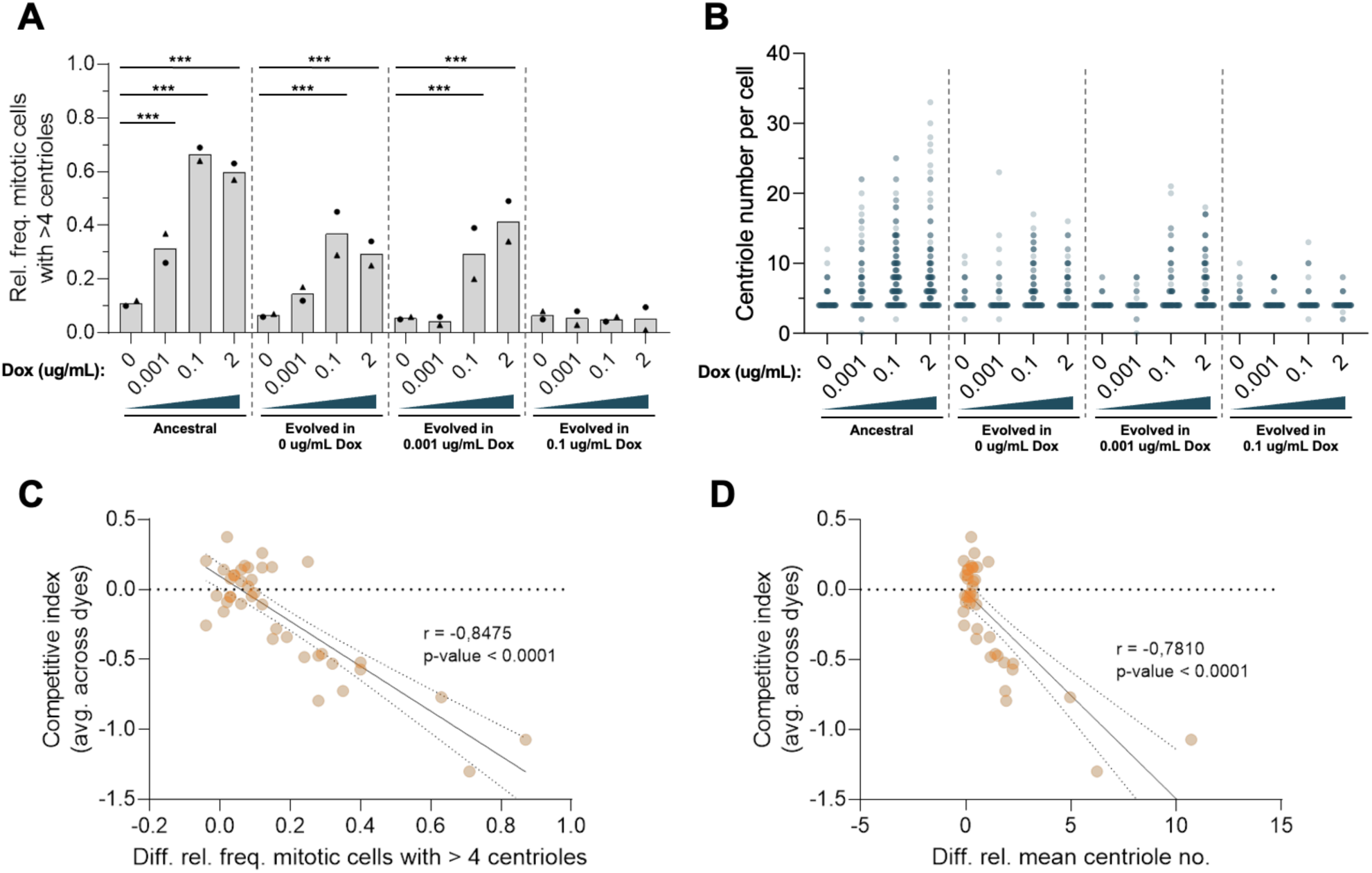
The fitness cost in ancestral and evolved populations is correlated with the level of centriole amplification. **A -** Relative frequency of mitotic cells with extra centrioles in ancestral and evolved MCF10A-PLK4 populations treated with Dox at the indicated concentrations for 24h. **B** - Centriole number distributions in ancestral and evolved MCF10A-PLK4 populations treated with Dox at the indicated concentrations for 24h. 95-100 cells were analysed for each condition in two independent experiments. Statistical analyses were performed using a generalised linear model with the presence/absence of extra centrioles in a cell as a binomial response variable and taking Dox concentration and population (ancestral, evolved in Dox0, evolved in Dox0.001, and evolved in Dox0.1) as independent categorical predictors, plus their interaction. Significant differences between the indicated condition and the one treated with Dox0 within the same population are represented with *** (*p-*value < 0.001). **C -** Correlation between average competitive index of the two dye combinations and the difference in the relative frequency of mitotic cells with more than four centrioles between co-cultured MCF10A-PLK4 and MCF10A-TetR. **D -** Correlation between average competitive index of the two dye combinations and the difference in mean centriole number in mitotic cells between co-cultured MCF10A-PLK4 and MCF10A-TetR. Data points include competitive indices calculated for three time points (at days 1-3, 31-33, and 64-66) and the corresponding centriole number data (at days 1, 31, and 64). The best fitting linear regression is indicated as a solid line. The dashed lines indicate the 95% confidence interval.

Together with the previous competition assays, these results indicate that fitness differences between co-cultured populations could be explained by the level of centriole amplification. Indeed, we observed a significant correlation between the relative frequency of mitotic cells with extra centrioles and average competitive index (Fig 4C). We also observed a significant correlation between mean centriole number and average competitive index, albeit slightly weaker (Fig 4D). These results show that the level of centriole amplification, in particular, the relative frequency of cells with extra centrioles, can explain most of the observed variation in the competitive index.

In summary, our results suggest that evolution of cells in Dox0.1 leads to a loss of the ability to amplify centrioles in response to the tested concentrations of Dox. This is not observed in the population evolved in Dox0.001, where cells do not amplify centrioles in low Dox but maintain that ability in higher Dox concentrations. These results suggest the existence of different regulatory mechanisms or the same mechanism operating at different magnitudes, that counteract centriole amplification at low Dox and high Dox.

### Slower proliferation explains the fitness cost of centriole amplification

Differences in competitive fitness can be due to changes in population growth. Centriole amplification can lead to defects in cell proliferation, cell cycle arrest and cell death^5,23^. Thus, we monitored the population size of the evolving populations by automated cell counting and calculated their relative growth rate compared with the Dox0-treated population. As expected, Dox treatment did not significantly affect the growth rate of control MC10A-TetR (Fig. S3A) and MCF10A-PLK4^1−608^ (Fig. S3B), when compared with Dox0 over time. In addition, we observed no significant change in the growth of MCF10A-PLK4 grown in Dox 0.001, whereas in Dox 0.1, the relative growth rate of the population decreased up to day 8 and progressively returned to basal levels (Fig. S3C). These results suggest that fitness differences can be attributed to changes in population growth.

Population growth depends on the balance between cell proliferation and death. We first assessed cell death in our competition assay setup by flow cytometry using propidium iodide staining, which labels dead or dying cells. The relative frequency of viable cells was not affected by Dox treatment in any of the cell lines, at any time point (Fig. 5A-B) suggesting that the fitness cost in induced MCF10A-PLK4 populations cannot be explained by differences in cell death.

**Figure 5:**
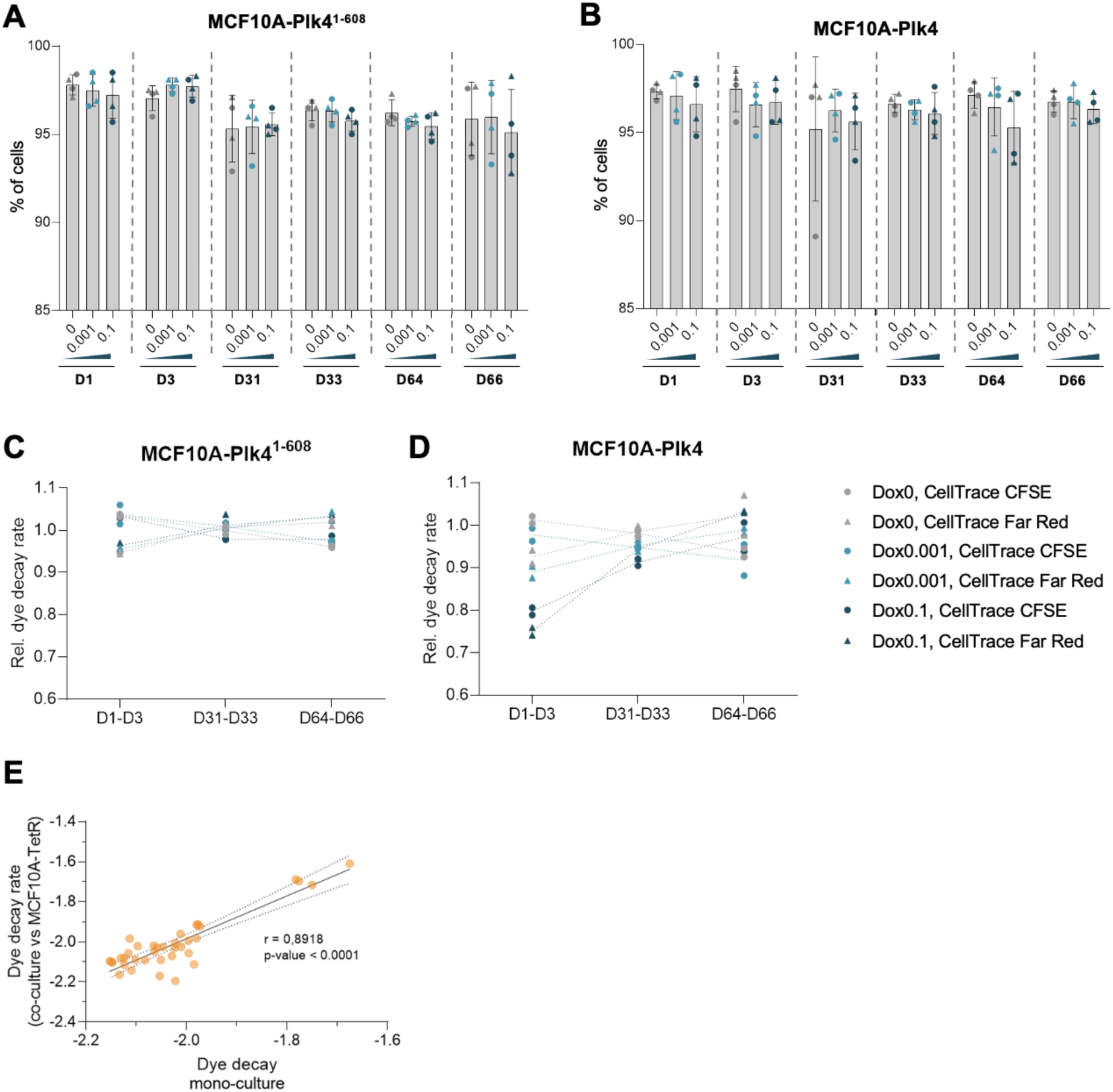
Slower proliferation explains the fitness cost of centriole amplification. **A, B -** Percentage of PI-negative MCF10A-PLK4^1–608^ (A) or MCF10A-PLK4 (B) co-cultured with MCF10A-TetR at the indicated time points. Data corresponding to co-cultures are represented as circles (MCF10A-PLK4^1–608^/MCF10A-PLK4 labelled with CellTrace CFSE) or triangles (MCF10A-PLK4^1–608^/MCF10A-PLK4 labelled with CellTrace Far Red). **C, D -** Relative dye decay rate of MCF10A-PLK4^1–608^ (C) or MCF10A-PLK4 (D) co-cultured with MCF10A-TetR at the indicated time points. The relative dye decay rate was calculated as a proxy for the rate of cell division. Data corresponding to co-cultures grown in Dox0 (light grey), Dox0.001 (blue), or Dox0.1 (dark blue) are represented as circles (MCF10A-PLK4^1–608^/MCF10A-PLK4 labelled with CellTrace CFSE) or triangles (MCF10A-PLK4^1–608^/MCF10A-PLK4 labelled with CellTrace Far Red). The dashed lines connect the average relative dye decay rate for each Dox concentration and dye combination in two independent experiments. See table S3 and S4 for statistical analyses. **E** - Correlation between dye decay rates of co-cultured (with MCF10A-TetR) and mono-cultured MCF10A-PLK4 populations at D1-3, D31-33, and D64-66. Data are represented as circles (MCF10A-PLK4 labelled with CellTrace CFSE) or triangles (MCF10A-PLK4 labelled with CellTrace Far Red). The best fitting linear regression is indicated as a solid line. The dashed lines indicate the 95% confidence interval.

Secondly, we assessed cell proliferation by measuring the decay rates of CellTrace CFSE and CellTrace Far Red in our competition assay setup. After labelling, each daughter cell inherits, on average, half of the dye molecules present in the mother cell following mitosis. Thus, the rate of dye decay can be used as a proxy for the rate of cell division.

As a control, we found no effect of the proliferation dye or initial frequency of each population on relative dye decay rates. Moreover, the two replicates were well correlated, validating the methodology (Fig. S4). As expected, control MCF10A-PLK4^1–608^ showed no significant differences in relative dye decay at any time point, regardless of Dox treatment (Fig. 5C). At days 1-3, the dye decay rate of induced MCF10A-PLK4 was significantly lower than MCF10A-TetR and was more affected by higher Dox concentration. This result suggests slower proliferation of cell populations with higher levels of centriole amplification (Fig. 5D). Importantly, over time, the relative dye decay rate of Dox0.001 and Dox0.1-treated MCF10A-PLK4 recovered at days 31-33, at which point only the population evolved in Dox0.1 showed significantly reduced decay rates. At days 64-66 (Fig. 5D) the relative dye decay rate of both Dox-treated populations was equal to MCF10A-TetR (relative dye decay close to 1). These results suggest that proliferation is the main factor impinging on fitness.

Several papers have now reported a potential non-cell-autonomous effect of centriole amplification. It was suggested that cells with centriole amplification secrete factors that promote invasion in neighbouring cells that do not show centriole amplification, thus contributing to changes in the microenvironment^9,23^. It is thus possible that cell proliferation is also sensitive to cell non-autonomous effects in the co-cultures. We tested this by performing mono-cultures and assessing for cell proliferation and comparing with the results in the co-cultures (Fig. 5E). We observed a good correlation between co-cultured and mono-cultured populations (Fig 5E), suggesting that potential non-cell-autonomous interactions between cells with and without centriole amplification did not affect their proliferative capacity.

### The fitness cost of centriole amplification is correlated with G1 delay and p53 activation

Changes in proliferation resulting from centriole amplification were previously associated with cell cycle delays resulting from p53 activation^6,53–55^. We next tested whether the changes in fitness observed were associated with those alterations.

We analysed the cell cycle profiles, by staining cells with Hoechst 33342, in our competition assay setup. As expected, control MCF10A-TetR and MCF10A-PLK4^1−608^ displayed no changes in their cell cycle profiles over time and for all Dox concentrations, either in mono-culture or in co-culture (Fig. 6A, Fig. S5). However, in MCF10A-PLK4 populations induced with Dox0.1 we observed that G1 cells were significantly overrepresented at day 3, both in mono-cultures and co-cultures (Fig. 6B, Fig. S5). Conversely, no significant differences were observed in the cell cycle profiles at days 31, 33 and 64, 66 consistent with the increase in fitness and dye decay rates observed at that time point, apart from a small reduction in the percentage of G1 cells at day 66 in the Dox0.1-treated population (Fig. 6B). No significant differences were observed in Dox0.001, in accordance with the reduced fitness costs observed for this condition. In summary, our data suggest that the initial fitness and proliferation deficit in Dox0.1 induced MCF10A-PLK4 populations results from a delay/arrest in G1.

**Figure 6:**
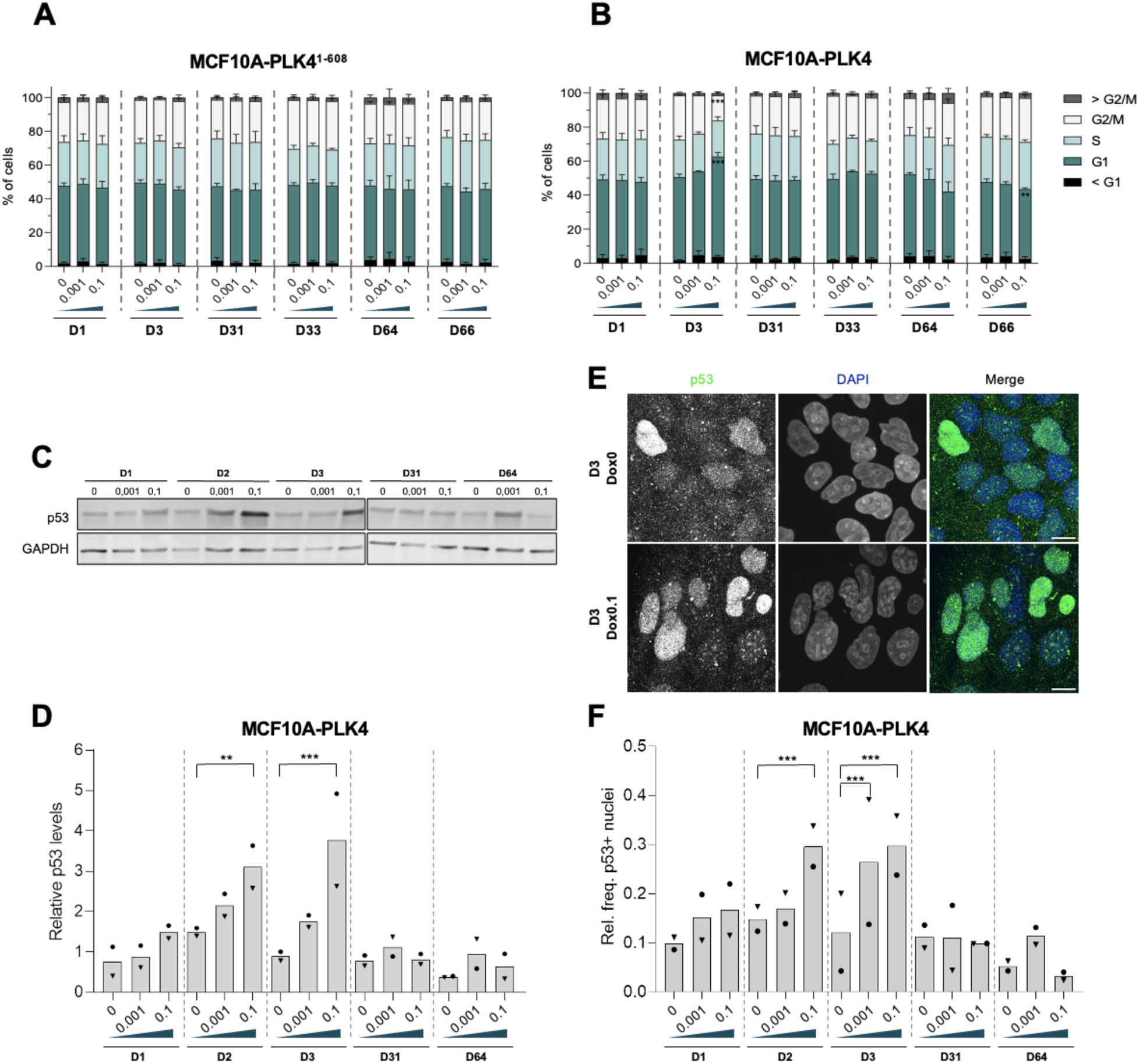
Cell populations with centriole amplification showed cell cycle defects and p53 activation. **A, B -** Relative frequency of cells in G1 (dark green), S (light green), and G2/M (white) for MCF10A-PLK4 (A) and MCF10A-PLK4 (B) in co-cultures at the indicated time points. Average relative frequencies of co-cultured populations in the respective cell cycle stage. The error bars represent the mean +- standard deviation. **C -** Representative immunoblot for p53 in MCF10A-PLK4 populations grown with the indicated Dox concentration, at the indicated time points. **D -** Quantification of p53 relative to GAPDH by immunoblotting in MCF10A-PLK4 populations treated with the indicated dose of Dox, at the indicated time points. **E -** Representative images of interphase cells containing p53-negative and positive nuclei. **F -** Relative frequency of p53-positive nuclei in MCF10A-PLK4 populations treated with the indicated dose of Dox, at the indicated time points. The bars indicate the average of two independent experiments. Symbols in D and F represent independent experiments. Statistical analyses in D and F were performed using a linear model with relative p53 intensity (by immunoblotting or by immunofluorescence) in a cell as a response variable and taking Dox concentration and time point as independent categorical predictors, plus their interaction. Significant differences between the indicated condition and the one treated with Dox0 at the same time point are indicated with *** (*p-*value < 0.001).

We next assessed p53 activation in our setup in the MCF10A-PLK4 cell line at different time points and levels of Dox. Activation of p53 is usually associated with an increase in its abundance and nuclear translocation^56^. We analysed p53 levels by western blot (Fig. 6C, D) and p53 nuclear localisation by immunofluorescence as a proxy for its activation (Fig. 6E, F). Our western blot data showed an increase in p53 levels that was higher at high Dox concentration in the first three days of treatment compared to non-induced MCF10A-PLK4. Whereas this change was also observed at Dox0.001 it was faster in Dox0.1, starting on day 1 and was more pronounced in day 2 and 3 (Fig. 3C, D). At days 31 and 64, when we observed no effects in cell cycle (Fig. 6B) or proliferation (Fig. 5D), p53 levels in induced MCF10A-PLK4 at all Dox concentrations were similar to those of the non-induced population (Fig. 6C, D).

Consistently with total protein levels, we observed an increase in the relative frequency of interphase cells with nuclear p53 over the first three days of induction, which correlated positively with the concentration of Dox. At days 31 and 64, p53 nuclear localisation reverted to basal levels despite continuous Dox treatment (Fig. 6F). Thus, p53 was activated over the first three days of induction, correlating with the observed cell cycle arrest/delay and the strongest fitness cost, and its activation decreased as the populations adapted. Interestingly, the results show that both p53 activation and cell cycle changes recovered more rapidly than population fitness, which was still significantly reduced at D31-33, suggesting that other factors may have contributed to the observed fitness differences. In conclusion, we observed short term proliferation defects, p53 activation and cell cycle delay/arrest following chronic PLK4 overexpression, as previously observed^6,10^, but these defects gradually disappeared over time, as the levels of centriole amplification decreased. This supports the idea that adaptation to long term induction of PLK4 overexpression occurred by suppression of centriole overproduction. We next tested potential molecular mechanisms associated with this adaptation.

### Centrosomal PLK4 levels were restored in evolved MCF10A-PLK4 populations despite its mRNA levels remaining elevated

In the previous sections, we dissected the processes underlying the fitness cost correlated with increased centriole numbers. This could have occurred by inhibition or loss of PLK4 overexpression, for example through loss of the transgene or hypermethylation of the promoter^57^, or inhibition of centriole overproduction.

We asked if the reduction in centriole amplifications could be explained by loss/down-regulation of PLK4 exogenous and/or endogenous expression. We first quantified PLK4 exogenous and endogenous expression by RT-qPCR. In MCF10A-PLK4^1–608^ populations, the PLK4 transgene was consistently overexpressed (Fig. S7). For MCF10A-PLK4 populations treated with Dox0.001, the levels of the exogenous PLK4 mRNA were not significantly elevated across the experiments (Fig. 7A). This is comparable to previous studies, which showed that small increases in PLK4 mRNA lead to centriole amplification^19,31,58^. Intriguingly, in MCF10A-PLK4 Dox 0.1-treated populations, we observed significant overexpression of the PLK4 transgene at days 1, 3, 46, and 64, but not at days 16 and 31 (Fig. 7A). Then, we measured the expression levels of the endogenous PLK4 gene. Expression levels were not altered in any of the cell lines, Dox concentrations and time points with exception of a decrease in endogenous PLK4 expression in the Dox 0.1 MCF10A-PLK4 population at day 3 (Fig. 7B, S7). We validated these results with a different set of primers targeting the exogenous or both the exogenous and endogenous PLK4 genes, and observed similar results (Fig S7). Thus, the PLK4 transgene was significantly overexpressed at Dox0.1, despite an apparent decrease at days 16 and 31. This suggests that transcriptional silencing may have contributed to the decrease in relative frequency of mitotic cells with extra centrioles at those stages. However, this cannot explain the loss in the capacity of evolved MCF10A-PLK4 populations in Dox0.1 to amplify centrioles at days 46 and 64 (Fig. 2C).

**Figure 7:**
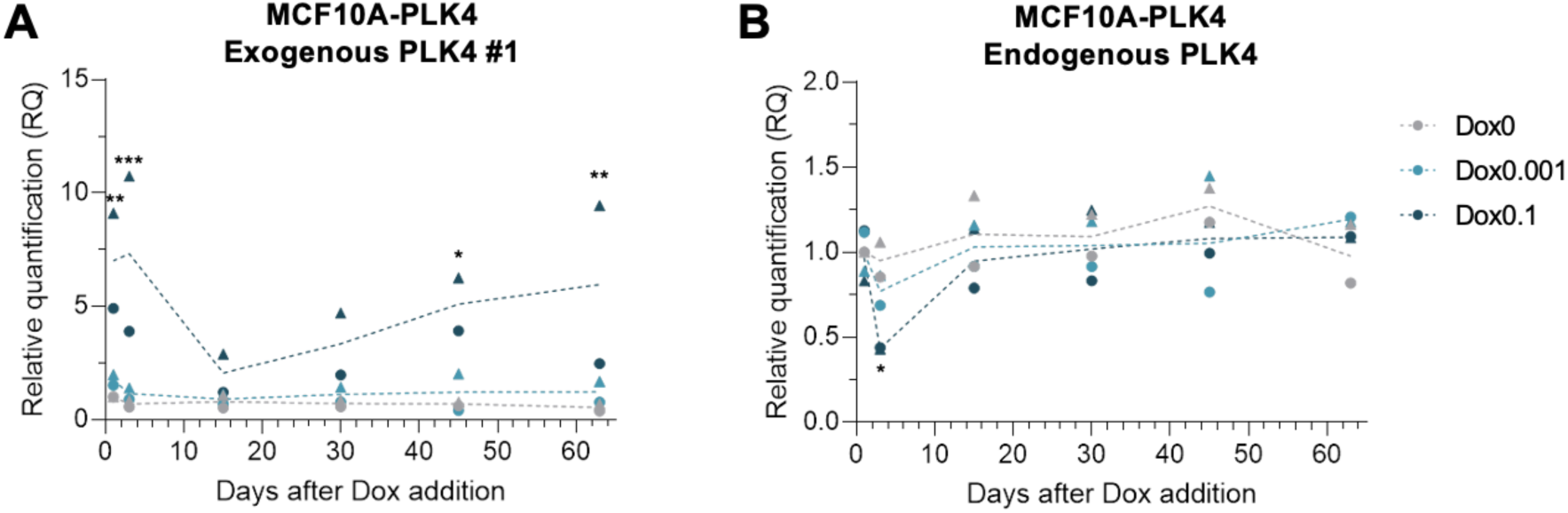
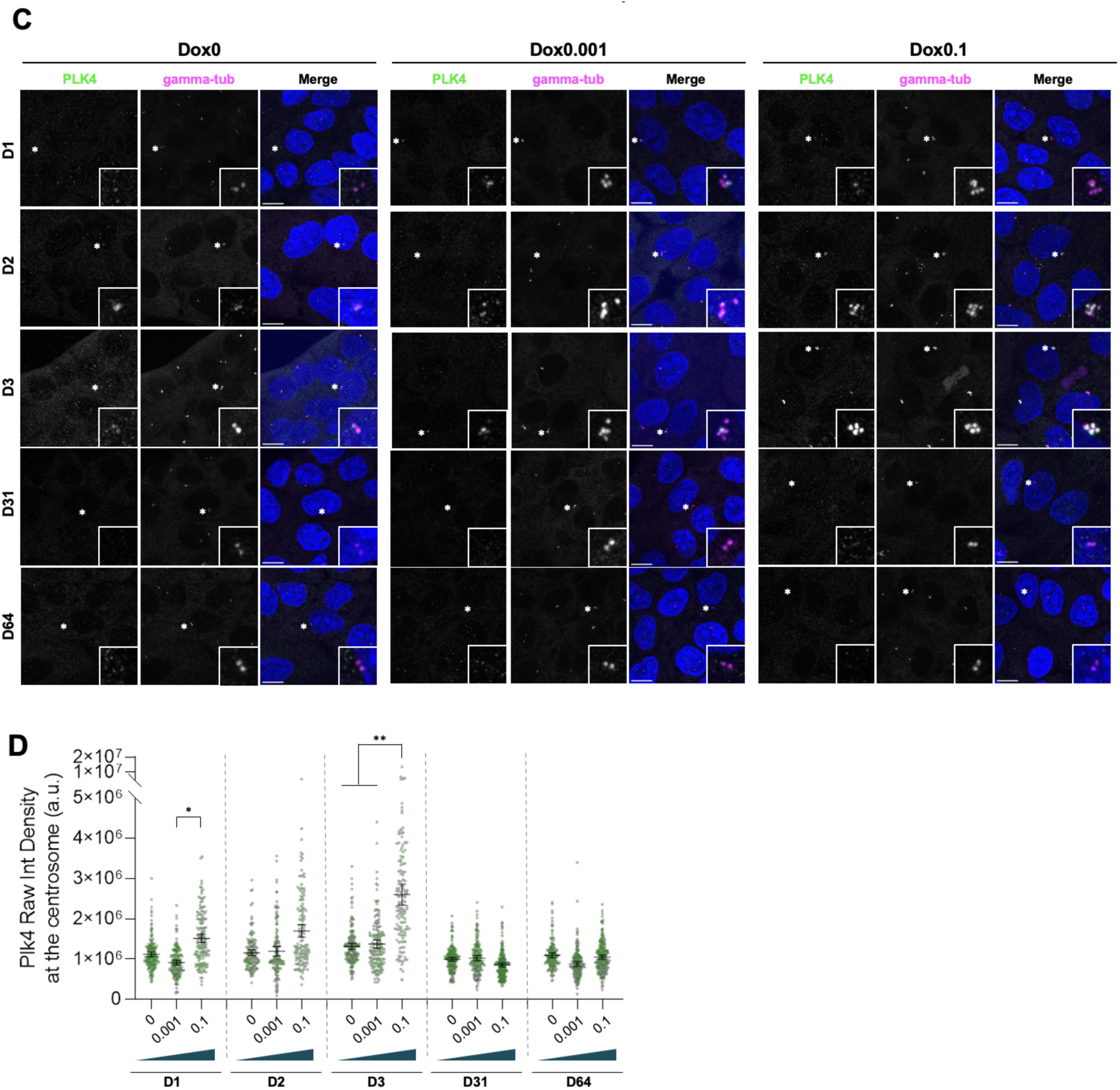
Centrosomal PLK4 levels were restored in evolved MCF10A-PLK4 despite its mRNA levels remaining elevated. **A, B -** PLK4 mRNA levels analysed by RT-qPCR. The primer sets used are specific for exogenous (A) and endogenous PLK4 (B). Dashed line corresponds to the average RQ in two independent experiments measured in RNA extracted from MCF10A-PLK4 populations grown with Dox0 (grey), Dox0.001 (blue), Dox0.1 (dark blue) μg/mL at the indicated time points. Statistical analyses were performed using linear mixed models taking Dox concentration and day as independent predictors, plus their interaction, and experiment as a random effect, with RQ as a response variable. **C -** Representative image of cells stained for PLK4 (green), gamma-tubulin (magenta) and DNA (blue). **D -** Levels of centrosomal PLK4 in MCF10A-PLK4 populations grown in Dox0, Dox0.001, and Dox0.1 at the indicated time points. 50 to 100 cells were quantified per condition for each of the two independent experiments. Colours (grey and green) represent independent experiments. Statistical analyses were performed using generalised linear models taking Dox concentration and day as independent predictors, plus their interaction, with the natural logarithm of the mean raw integrated density as a response variable. Significant differences between the indicated conditions in A, B, and D and the Dox-0 control at the respective day are represented with * (*p*-value<0.05), ** (*p*-value<0.01), and *** (*p*-value<0.001).

STIL and SAS-6 are two key proteins for proper centriole assembly and their recruitment to the centrosome is dependent on PLK4^29,30,59–61^. Therefore, we asked if downregulation of STIL or SAS-6 could be compensating for PLK4 overexpression. However, we did not observe any significant differences in the expression levels of STIL or SAS-6 in induced versus non-induced MCF10A-PLK4 at any time point (Fig. S8). Thus, the reduction of centriole amplification observed could not be explained by down-regulation of these genes.

Despite PLK4 remaining overexpressed, other regulatory mechanisms could be preventing centriole overproduction, such as inhibition of translation, increase in protein degradation, decrease in kinase activity, or impaired centrosomal recruitment. Thus, we performed immunostaining of PLK4 and measured its levels specifically at the centrosome by immunostaining. For MCF10A-PLK4 populations in Dox0.1 we observed an increase in the centrosomal PLK4 over the first three days, which returned to basal levels at days 31 and 64 (Fig. 7C,D). At days 31 and 64 there were no significant differences between populations grown and the corresponding non-induced populations (Fig. 4D). Thus, PLK4 overexpression levels as measured by RT-qPCR were correlated with an accumulation of centrosomal PLK4 at the first time points. However elevated PLK4 mRNA did not translate into higher levels of PLK4 at the centrosome in the later stages of the experiments. These results suggest that the long-term suppression of centriole amplification could be explained by a reduction in the levels of centrosomal PLK4. Thus, whereas initially centriole amplification is countered by reduced proliferation of cells with extra centrioles and down-regulation of PLK4 overexpression, at later stages, a different mechanism may block PLK4 biosynthesis or centrosomal recruitment irrespective of high RNA levels.

### Suppression of centriole amplification in evolved MCF10A-PLK4 populations treated with Dox is irreversible

We showed that loss of centriole amplification was correlated with an increase in relative fitness and reduction in the levels of centrosomal PLK4 despite its continued overexpression, suggesting a long-term inhibition of the production of extra centrioles in MCF10A-PLK4 evolved in Dox0.1 at all Dox levels and in MCF10A-PLK4 evolved in Dox0.001 in 0.001 Dox. This suggests some degree of irreversibility of evolution. So, we next asked if this reduction in centriole amplification capacity was reversible. We cultured evolved MCF10A-PLK4 populations without Dox for 20 days. Then, we treated them with different concentrations of Dox for 24h and analysed centriole numbers. We observed that cell populations that evolved without Dox experienced a dose-dependent increase in the relative frequency of mitotic cells with extra centrioles when they were treated with Dox at 0.001 or 0.1 μg/mL (Fig. 8A Left). In contrast, evolved populations grown with Dox0.001 or 0.1 μg/mL did not show an increase in centriole amplification when treated with those concentrations, respectively (Fig. 8A, Center and Right). Similarly, when as a control, these populations were maintained at the respective concentration of Dox, we observed no significant differences in the relative frequency of mitotic cells with extra centrioles.

**Figure 8:**
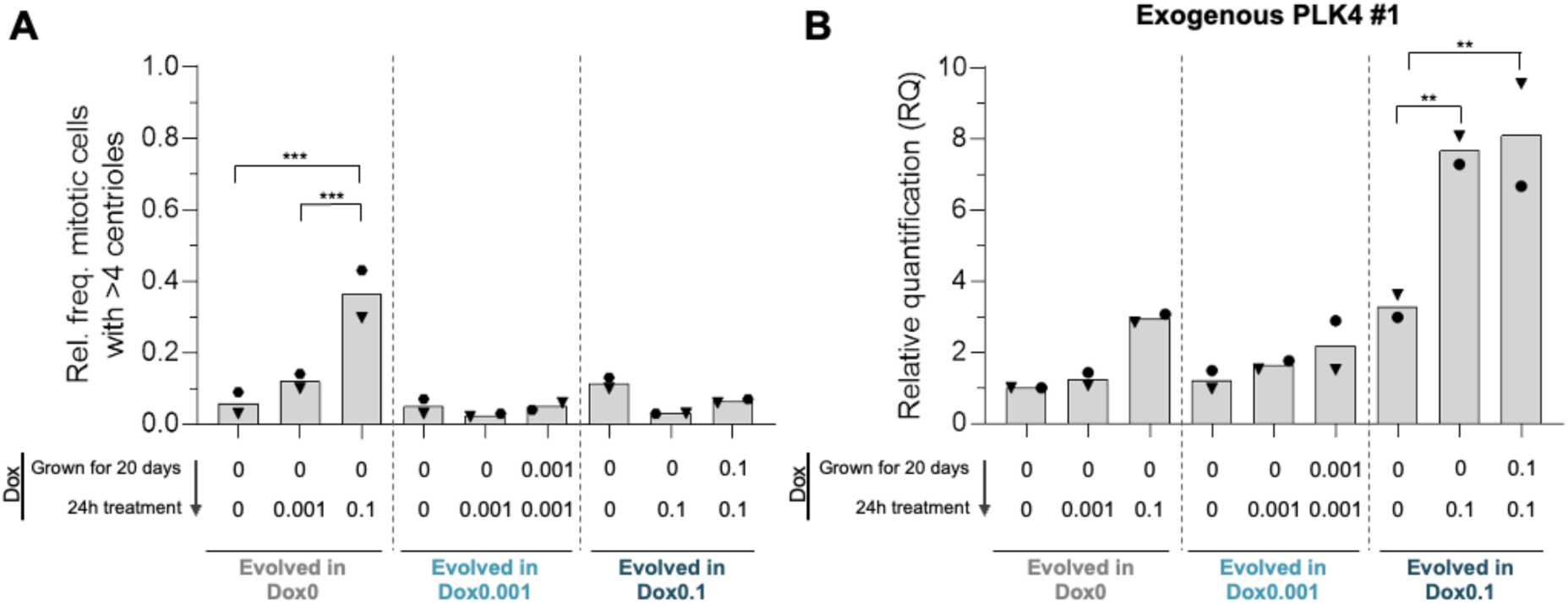
Suppression of centriole amplification in evolved MCF10A-PLK4 populations treated with Dox is irreversible. **A** - Relative frequency of mitotic cells with more than four centrioles in populations evolved in 0, 0.001 or 0.1 μg/mL of Dox, grown for additional 20 days without Dox and treated with the indicated doses of Dox for 24h. As controls, cells were grown in parallel with the same dose of Dox as during experimental evolution. At least 50 cells were analysed per condition for each of the two independent experiments. Symbols represent independent experiments. **B** - PLK4 mRNA levels analysed by RT-qPCR. The primer sets used are specific for exogenous (PLK4 (B). Dashed line corresponds to the average RQ in two independent experiments measured in RNA extracted from MCF10A-PLK4 populations grown with Dox at 0 (grey), 0.001 (blue), and 0.1 (dark blue) μg/mL at the indicated time points.

Therefore, the repression of centriole amplification in these populations was stable over time despite induction having been relieved. We concluded that chronic Dox treatment induced a potentially irreversible loss in the capacity of MCF10A-PLK4 to generate extra centrioles. Finally, we analysed the level of PLK4 overexpression in these populations (Fig 8B). Treatment with Dox0 and Dox0.001 did not significantly affect the expression of the PLK4 transgene in any condition. In contrast, treatment with Dox0.1 induced significant overexpression of the PLK4 transgene in Dox0.1-evolved populations grown for 20 days with or without Dox, without resulting in centriole amplification. Intriguingly, the Dox0-evolved population grown for 20 days without Dox and treated with Dox0.1 showed a non-significant increase in the expression level of the PLK4 transgene. We observed similar trends of PLK4 expression using a different primer set for the transgenic mRNA and primers targeting both the exogenous and endogenous PLK4 transcripts. These results suggest that evolved populations were generally less sensitive to induction of PLK4 overexpression by Dox. Moreover, these results might explain the less extreme levels of centriole amplification in Dox0-evolved populations when treated with Dox compared with the ancestral population.

### Evolutionary dynamics upon PLK4 overexpression can be partially explained by the presence of low-frequency MCF10A-PLK4^1–608^ in the ancestral population

The observed evolutionary dynamics could potentially be explained by cells bearing centriole amplification being outcompeted by a mutant already present in the ancestral population that could not generate extra centrioles, or by the appearance of a mutant during the evolution experiment. We addressed this hypothesis by performing bulk RNA-seq of cells sampled at days 1, 3, 31, and 64 (final time point) of experimental evolution.

We started by performing variant analysis of PLK4 to assess the presence of mutations in this molecule that could inhibit centriole overproduction. If this hypothesis was correct, we expected that such a variant would be observed at low frequencies or not detected in the ancestral population and rise in frequency over time. As expected, we did not detect variants displaying such a pattern in MCF10A-TetR and MCF10A-PLK4^1–608^. For MCF10A-PLK4, we detected reads containing a mutation (A to T) which results in a stop codon yielding the truncation of the exogenous PLK4 present in MCF10A-PLK4^1–608^ (Fig. 9A, B). Additionally, we detected a synonymous SNP (G to A) that is also present in MCF10A-PLK4^1–608^ (Fig. 9C, D). We detected both these SNPs at days 31 and day 64 in both evolution experiments for the populations treated with Dox 0.1, being higher in the latter time point. The SNP was not detected in MCF10A-PLK4 evolved in Dox0 or Dox0.001. These results suggest that the ancestral MCF10A-PLK4 population was contaminated with a very small proportion of MCF10A-PLK4^1–608^ and that with higher Dox concentration imposing a stronger selective pressure against cells with centrosome amplification, the cells overexpressing the truncated version of Plk4 became more abundant over time.

**Figure 9.**
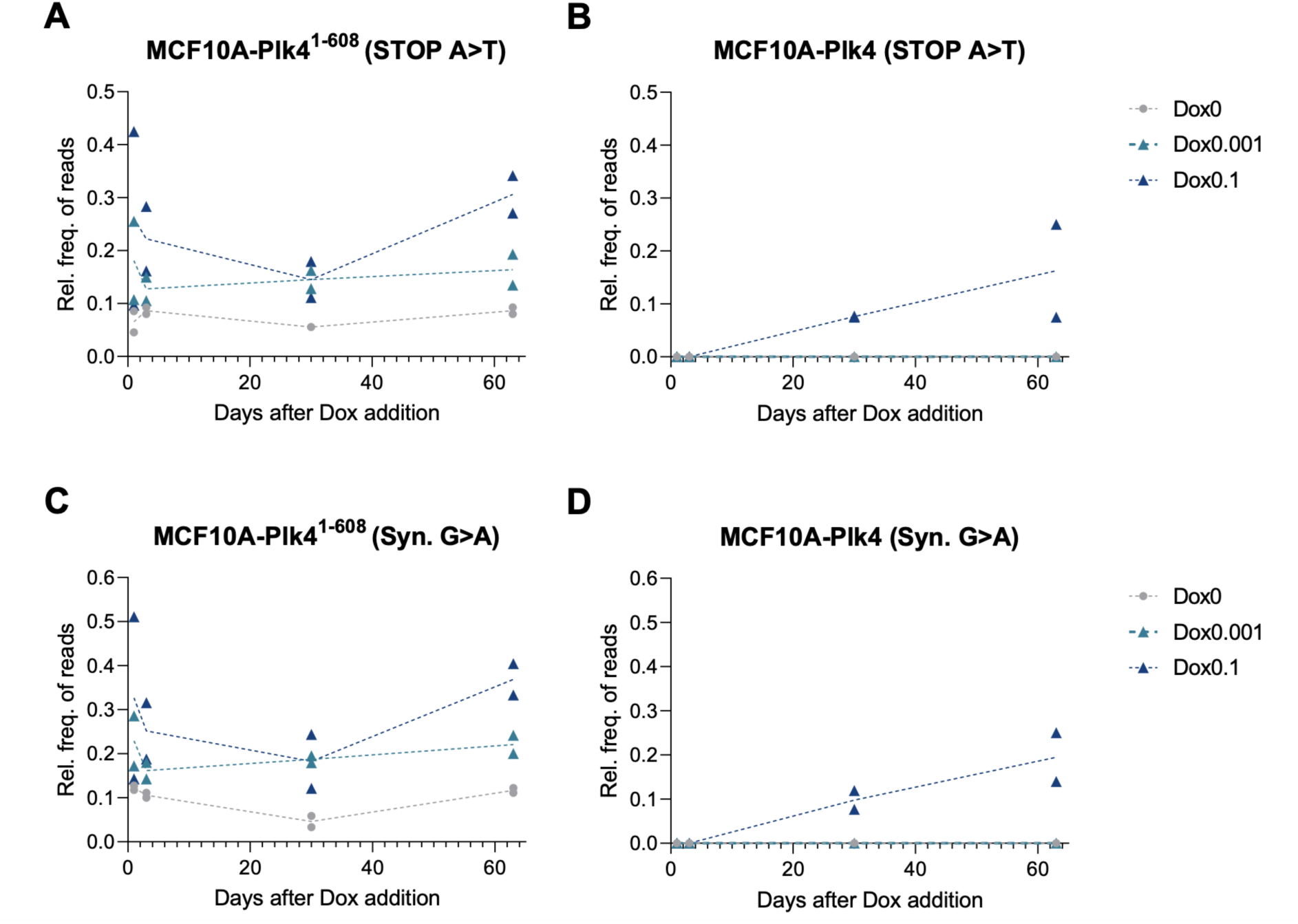
MCF10A-PLK4^1–608^ specific SNPs relative frequency increased in the MCF10A-PLK4 samples over the time course of the experiment. Relative frequency of RNA-seq reads with the indicated SNP leading to a STOP codon (A and B) or to a synonymous SNP (C and D) for each time point after dox addition, and Dox concentration (Dox0 in grey, Dox0.001 in light blue and Dox0.1 in dark blue). A and C refer to MCF10A-PLK4^1–608^ samples, while B and D refer to MCF10A-PLK4 samples. Results correspond to two independent experiments.

### MCF10A-PLK4 appear to converge to centriole overproduction-selection balance upon PLK4 overexpression

Since our interpretation of the results was confounded by the contamination of MCF10A-PLK4 populations with MCF10A-PLK4^1–608^, we resorted to repeating all the experiments with a different batch of ancestral MCF10A-PLK4. As with the original experiments, we observed an initial increase of centriole amplification proportional to the dose of doxycycline (Fig. 10A). In these new experiments, the average relative frequency of cells with extra centrioles in populations treated with Dox0.001 was 52,5% and in populations treated with Dox0.1 was 86% (Fig. 10A). Whereas the level of centriole amplification progressively decreased both in Dox 0.001 and Dox 0.1 at day 3 and day 31, it was not significantly altered between day 31 and day 64 (Fig. 10A). Strikingly, the relative frequency of cells with extra centrioles was indistinguishable between MCF10A-PLK4 populations evolved in Dox 0.001 and Dox 0.1 at day 64. Consistently, the fitness deficit of these populations relative to MCF10A-TetR was proportional to the degree of centriole amplification: initially, MCF10A-Plk4 treated with Dox0.1 experienced a stronger fitness penalty, that was eventually ameliorated, whereas the fitness penalty of MCF10A-PLK4 treated with Dox0.001 remained approximately constant throughout experimental evolution (Fig. 10B). As previously, we observed that the fitness penalty was correlated with cell cycle delay/arrest (Fig. 10C). Together, these results suggest that MCF10A-PLK4 eventually converged to similar centriole amplification and relative fitness set points regardless of the level of PLK4 overexpression.

**Figure 10.**
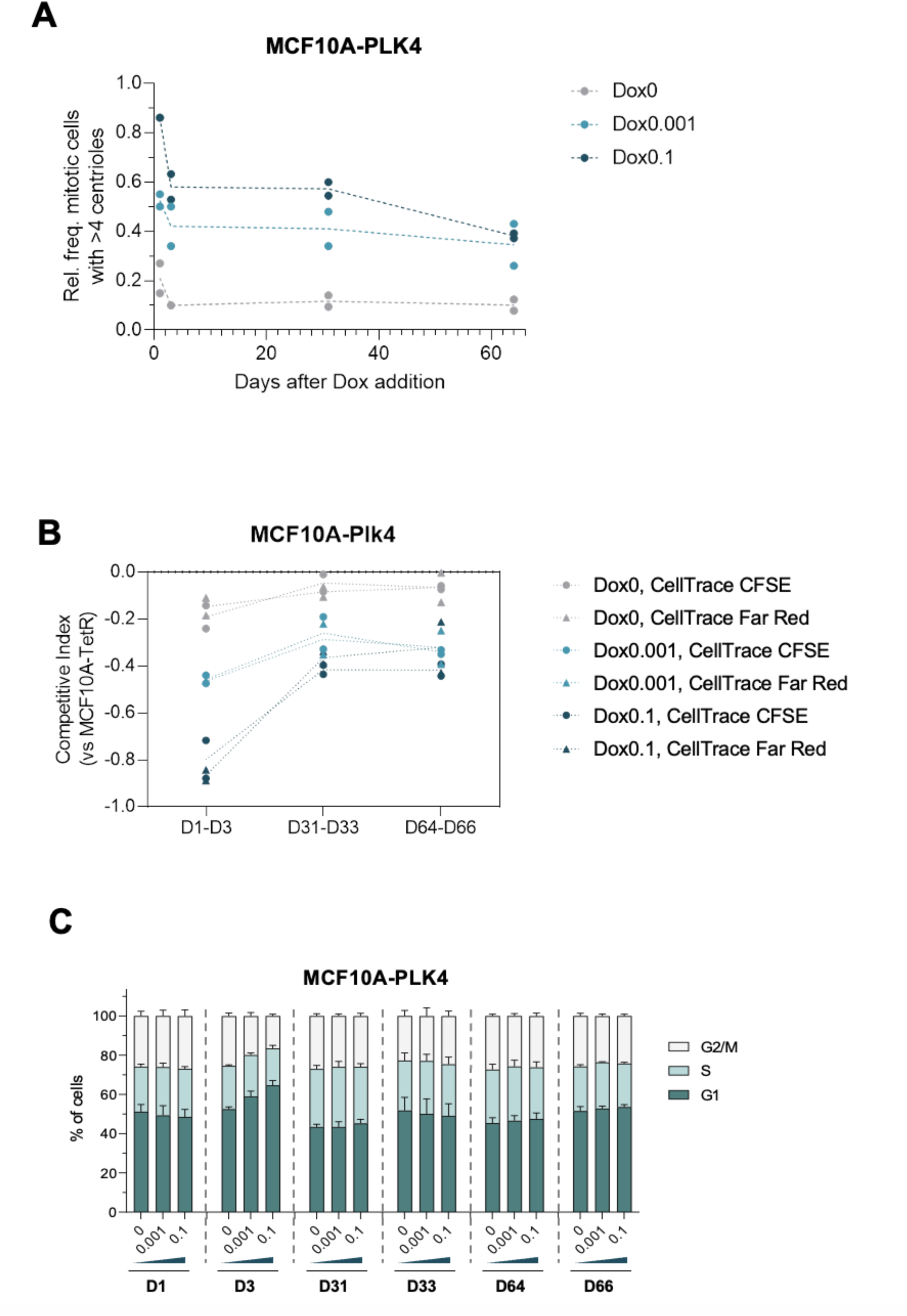
MCF10A-Plk4 induced with different doses of Dox seem to converge to a new equilibrium of centriole amplification levels. MCF10A-Plk4 cells were cultured with different doses of doxycycline (Dox0, Dox 0.001or Dox 0.1) for 2 months. (A) Relative frequency of mitotic cells with extra centrioles (more than four) at each time point for populations grown in Dox0 (grey), Dox0.001 (light blue), or Dox0.1 (dark blue). The dashed line represents the mean of the two independent experiments. (B) Competitive index of MCF10A-PLK4 relative to MCF10A-TetR at the indicated time points. Note that for each experimental situation there are 4 data points (independent experiments and swapping of the dye). The dashed lines connect the average competitive index corresponding to each Dox concentration and dye combination in two independent experiments. (C) Relative frequency of MCF10A-PLK4 cells in G1 (dark green), S (light green), and G2/M (light grey) at the indicated time points. Average relative frequencies of co-cultured populations in the respective cell cycle stage from the two independent experiments. For each cell cycle phase and each time point, the log odds ratio was calculated for each phase against the other two. A linear model was fitted with Dox concentration as a categorical predictor. Error bars represent the mean +- standard deviation.

## DISCUSSION

In this study, we conducted the first systematic assessment of how cell populations evolve in response to different levels of centriole amplification induced by chronic PLK4 overexpression. More specifically, we quantified fitness, cell proliferation, centriole numbers, and PLK4 dynamics and how these parameters correlate and change over time. However, we identified the presence of MCF10A-PLK4^1–608^ cells in MCF10A-PLK4 populations in our original experiments. Specifically, our results suggested that induced MCF10A-PLK4 populations suffered a fitness penalty proportional to the level of centriole amplification. Thus, these cells were probably outcompeted by the initially low-frequency MCF10A-PLK4^1–608^, in which induction does not lead to centriole amplification^64^. It is noteworthy that the population evolved in Dox0.001 maintained some centriole amplification capacity upon exposure to increased Dox doses. This observation may be due to a slower increase in frequency of MCF10A-PLK4^1–608^ in the population, compared with the Dox0.1 condition, due to the weaker selective pressure. However, we cannot exclude the possibility that this observation is caused by a cellular response of the MCF10A-PLK4 in the population, that somehow inhibited the formation of supernumerary centrioles. Interestingly, our experiments with uncontaminated MCF10A-PLK4 showed cells treated with Dox0.001 or Dox0.1 both converged to the same level of centriole amplification, irrespective of doxycycline concentration. These populations displayed fitness penalties along experimental evolution which similarly converged and stabilised, suggesting physiological adaptation to centriole amplification. Further work i exploring PLK4 expression dynamics in these populations, the basis for fitness differences, and the molecular causes of adaptation will elucidate the mechanisms that allow centriole amplification in cell populations.

Whereas in our original experiments the presence of MCF10A-PLK4^1–608^ could partially explain the desensitization of evolved MCF10A-PLK4 to overexpression induction, in our repeated experiments the evolved populations treated with Dox were still permissive to centriole amplification. It will be interesting to challenge them with different doxycycline concentrations and also test if the partial reduction in centriole amplification levels is reversible or not. Whereas in the first case, irreversibility can be attributed to MCF10A-PLK4^1–608^ taking over the population, it is possible that different dynamics are observed in uncontaminated MCF10A-PLK4. For example, if the reduction in centriole amplification levels is reversible, it will also be interesting to understand if the reversibility occurs in the same time frame for both populations evolved in the different Dox concentrations.

Our results also reveal potentially interesting nuances concerning centriole overproduction-selection balance. This hypothesis states that cells with centrosome amplification reach some equilibrium frequency depending on the rate of centriole overproduction and the strength of negative selection against cells with extra centrioles. For example, typical healthy cells experience few errors in centriole biogenesis (low overproduction rate) and strong negative selection, such that cells with centriole number anomalies are relatively rare. In cancer cells, in which errors are more common or compensatory mechanisms may exist, the level of centriole amplification tends to be higher. Under this premise, we expected that low levels of PLK4 overexpression (low, constant overproduction) would lead the population to a modest, stable level of centriole amplification, whereas high levels of PLK4 overexpression (high, constant overproduction) culminated in a higher fraction of the population with extra centrioles. The observation that both levels of PLK4 overexpression resulted in populations with similar degrees of centriole amplification suggests some cellular response that leads to a modulation of either centriole overproduction or selection. In the future, it will be interesting to perform RNA-seq or proteomics analysis to inquire about the molecular pathways that could underlie this adaptive response. In addition, these analyses may shed light on how centriole amplification is tolerated in cancer or may contribute to tumorigenesis. Nevertheless, it would be important to do similar studies in different cell types, including cells from different tissues to test if they respond in a similar fashion or if different tissues have different responses to prolonged induction of centriole amplification. This is particularly interesting because different tissues seem to have different tolerance to centriole amplification^65^, even though some of these response mechanisms may be conserved. Moreover, comparing the results obtained in different tissues with the amplification levels of tumors from matched tissues could show whether there is a correlation between both results, i.e., if there may be a tissue intrinsic tolerance that impacts on the level of centriole amplification of tumors.

## MATERIALS AND METHODS

### Cell culture

We cultured MCF10A-TetR, MCF10A-PLK41−608, and MCF10A-PLK4 as described previously^11^. We cultured the cells in DMEM (Gibco) and F-12 (Gibco) at 1:1 ratio, supplemented with horse serum at 5%, L-glutamine (Thermo Scientific) at 0.5 *μ*g/mL, penicillin and streptomycin (Thermo Scientific) at 100 U/mL, insulin (Sigma) at 10 *μ*g/mL, hydrocortisone (Sigma) at 100 ng/mL, cholera toxin (Sigma) at 1 *μ*g/mL, epidermal growth factor (Sigma) at 20 ng/mL, and maintained them at 37◦ C in a 5% CO2 atmosphere. All cell lines were tested for the presence of mycoplasma.

For the evolution experiment, we passaged cells every 72h and subcultured them in 150 mm dishes ensuring that cells were sub-confluent at the time of passaging, such that they predominantly spent time in exponential growth. The medium for doxcycyline-treated populations was further supplemented with doxycycline to a final concentration of 0.001 or 0.1 μg/mL. During passaging, cells were counted using a Scepter 2.0 Handheld Automated Cell Counter (Merck Millipore) and 60 μm Sensors (Merck Millipore). The growth rate of each condition was calculated taking the number of cells at passaging, Npass, and the initial seeding cell number, *N_seed_*, which we fixed at 2.2×106 cells, according to:

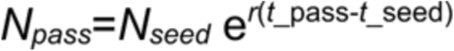

where *r* is the exponential growth rate (per day), *t*_pass is the day of passaging, and *t*_seed is the day when the population was seeded. Relative growth rates were then obtained by taking the growth rate for a given condition, at a given time point *t_pass*, and dividing it by the growth rate of the Dox0-treated population of the same cell line at the respective time point.

### Competition assays and flow cytometry

We seeded 1.5*x*10^6^ cells at days −1, 29, and 62 of each experimental evolution replicate in two T-25 flasks per condition, such that confluency was not reached in the next day and cells maintained exponential growth. At day 0 of each competition assay, CellTrace CFSE (Thermo Scientific) or CellTrace Far Red (Thermo Scientific) staining was performed according to manufacturer instructions. In brief, we replaced the cell culture medium in the flasks with 3 mL of 10 *μ*M CellTrace CFSE or 2 *μ*M CellTrace Far Red in PBS and incubated the cells at 37°C for 20 minutes. After staining, we washed the flasks with an equal volume of medium supplemented with 5% horse serum to remove unbound dye molecules. Then, we detached, centrifuged and resuspended the cells in the corresponding media. Co-cultures were prepared by mixing populations stained with CellTrace CFSE and CellTrace Far Red at 1:1 ratio. We seeded monocultures and co-cultures such that the confluency at the time of harvest was approximately 70-80%. At days 1 through to 3, cells were stained with 5 *μ*g/mL Hoechst 33342 (Invitrogen) in cell culture medium for 30 minutes at 37°C, before detaching them. Finally, we resuspended the cells in 1 *μ*g/mL propidium iodide (PI, Sigma) staining solution and incubated them at 37°C for 10 minutes. Mono- and/or co-cultures were then analysed by flow cytometry.

Flow cytometry analysis was conducted using a BD Fortessa X-20 cytometer and BD FACSDiva software. The cell population was gated based on their forward- and side-scatter area parameters (FSC-A and SSC-A, respectively) and doublets were excluded by plotting the area against the width of the forward-scatter parameter. Exclusion of dead cells was performed by gating out PI-positive cells. Finally, we gated live single-positive CellTrace CFSE and CellTrace FarRed cells for competition, proliferation analysis, and cell cycle analysis. Live cells in each cell cycle stage were manually gated. For viability analysis, we identified CellTrace CFSE- and CellTrace Far Red-positive singlets and excluded PI-positive cells in each sub-population. We acquired approximately 20,000 live cells for each sample during the evolution experiment and approximately 10,000 live cells for the remaining competition assays. Flow cytometry data was analysed using FlowJo.

### Quantification of competitive index, and dye decay rate

As mentioned in the previous section, we first gated single-positive CellTrace CFSE and CellTrace Far Red populations. We observed a small population of double-positive cells, as reported in the literature, which we excluded from further analysis. The relative frequency, of each sub-population*I*, *p_CFSE_* and *p_FarRed_*, at day *t* was calculated according to the following expressions:

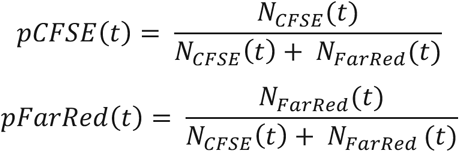

where *NCFSE*(*t*) is the total number of CellTrace CFSE single-positive cells, *NFarRed* the total number of CellTrace Far Red single-positive cells. As a proxy for fitness, we employed the competitive index, given by:

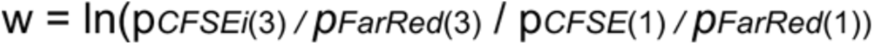

To calculate the mean rate of dye decay, as a proxy for the mean cell division rate, we first computed the average CFSE/Far Red intensity in single-positive cells at each day of the competition assay. As above, double-positive cells were excluded. As a simplification, we assumed constant dye decay; we estimated the mean dye decay rates for each cell line and Dox concentration according to:

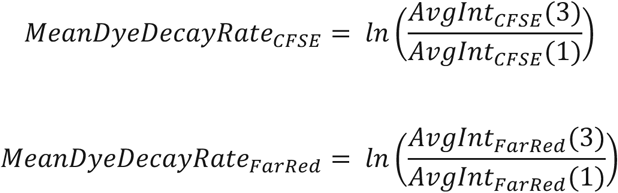

Relative dye decay rates were then calculated by dividing the mean dye decay rates of MCF10A-PLK4/MCF10A-PLK4^1–608^ by those of MCF10A-TetR in the same Dox conditions. Estimation of competitive index and mean dye decay rate was performed using R.

### Immunofluorescence and microscopy

For staining and imaging of centriolar proteins, cells were grown on 13 mm coverslips in 24-well plates, using the appropriate cell culture medium and fixed with 100% methanol at −20°C for 10 minutes. For immunostaining, we started by incubating cells with blocking solution (10% FBS in 1X PBS) for 30 minutes at room temperature. Samples were incubated with primary antibody dilutions (prepared in blocking solution) for 1 hour at room temperature or overnight at 4°C. Cells were then washed with 1X PBS and incubated with secondary antibody dilutions for 1 hour at room temperature. DNA staining was performed by incubating the cells with 1 *μ*g/mL Hoechst 33342 for 15 minutes at room temperature. Alternatively, cells were stained with DAPI during secondary antibody incubation. For immunostaining of p53, cells were incubated with a solution containing the p53 primary antibody and nanobody which were previously incubated together for 1 hour at room temperature, prepared according to manufacturer instructions. Finally, we mounted coverslips on microscopy slides, using Vectashield mounting medium (Vector Laboratories). Primary antibodies used: anti-CEP135 1:500 (rabbit, Abcam), anti-Centrin 1:500 (mouse, clone 20H5, Millipore); anti-PLK4 1:500 (rabbit, Metabion), anti-p53 1:500 (mouse, Millipore), anti-gamma-tubulin (mouse, clone GTU88, Sigma). Secondary antibodies used were: goat/donkey anti-mouse Alexa Fluor 488 1:500 (Molecular Probes), donkey anti-rabbit Rhodamine Red 1:500 (Jackson Immunoresearch), donkey anti-rat Alexa Fluor 647 (Molecular Probes). Nanobody (Atto 542 FluoTag-X2 anti-Mouse Kappa light chain, Nanotag). Images of mitotic cells for assessing centriole numbers and interphase cells for quantification of PLK4 levels and nuclear p53 were acquired using an Eclipse Ti-2 inverted microscope with 3i Marianas spinning-disk using a 100X objective and Andor Dragonfly spinning-disk, respectively.

### Image analysis

Microscopy images were analysed using Fiji/ImageJ. Centriole number counting was performed on 3D stacks. Each Centrin focus co-localising with a CEP135 focus was scored as a centriole. To assess centrosomal PLK4 levels, we applied a threshold (Triangle) in the gamma-tubulin channel to define ROI corresponding to the centrosome. Then, we used these ROI to measure PLK4 intensity (Raw Integrated Density) in its sum projections. To quantify presence/absence of nuclear p53, we adjusted the histograms of the p53 channel to a fixed range in order to compare between images and scored p53-positive nuclei.

### Western blotting

Cells lysis was performed by incubating cells in lysis buffer (Tris 10 mM pH7.4, EDTA 5 mM, NaCl 100 mM, Triton X-100 1%, Na3VO4 0,2 mM, NaF 50 mM, DTT 1 mM, Protease inhibitors (Roche)) on ice for 30 min. Then, samples were centrifuged at 13 500 rpm for 15 min at 4°C and the supernatant was transferred to a new tube. Protein extracts were quantified using the Bradford method and 60 *μ*g of total protein (per sample) were run and separated on sodium dodecyl sulphate-polyacrylamide gel electrophoresis (SDS-PAGE) and transferred onto nitrocellulose membranes. For western blotting, the membranes were blocked with 5% milk in TBS 1X, following incubation with the primary antibodies anti-p53 (1:500, mouse, (Ab-6) Pantropic OP43 Sigma-Aldrich) and anti-GAPDH (1:1000, rabbit, 14C10 Cell Signaling) for 1h at room temperature. Then, membranes were washed with TBS 1X, incubated with secondary antibodies IRDye 800CW Goat anti-Mouse (1:10 000, #926-32210, Li-cor) and IRDye 680RD Goat anti-Rabbit (1:10 000, #926-68071, Li-cor) for 1h at room temperature. Membranes were then washed and developed in Odyssey (Li-cor). Band intensity was quantified using Fiji/ImageJ.

### RNA extraction and RT-qPCR

RNA extraction was performed using the RNeasy Mini Kit (Qiagen) according to manufacturer instructions. For each sample, we performed on-column DNA digestion using RNase-free DNAse (Qiagen) as specified by the kit. RNA quantity and purity were assessed using NanoDrop 2000 (Thermo Scientific). We performed cDNA synthesis using the High Capacity RNA-to-cDNA kit (Applied Biosystems), according to manufacturer instructions.

For the RT-qPCR reactions we used iTaq SYBR Green (BioRad) according to kit instructions. Samples were prepared in triplicate in 384-well plates. The qPCR run and data analysis were performed using QuantStudio (Applied Biosystems). The following primers were used for quantifying the mRNA levels of GAPDH, exogenous PLK4 *truncated or full-length), endogenous PLK4, total PLK4 (i.e. primers anneal to both the exogenous and endogenous sequences), STIL, and SAS-6.

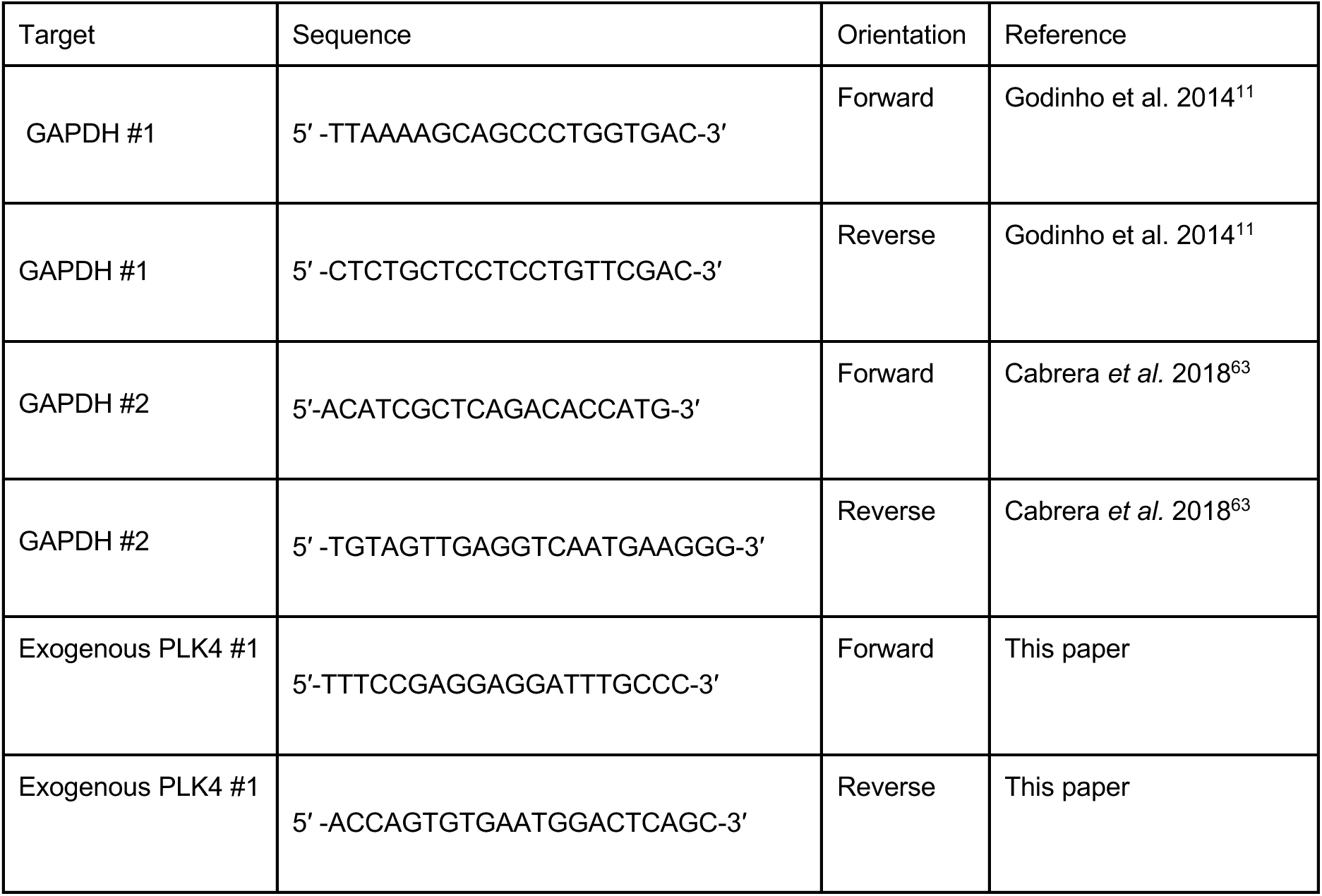
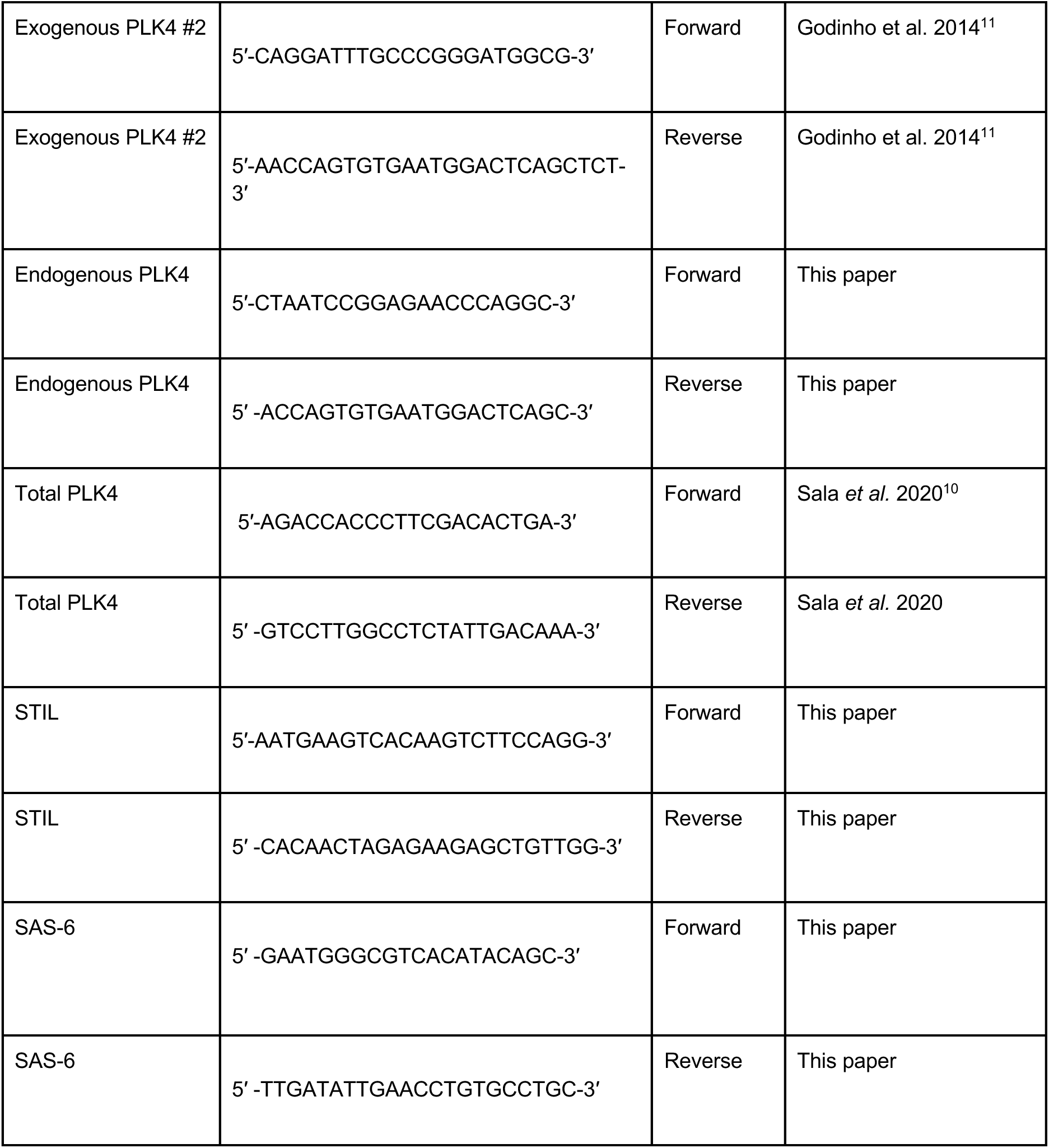

### Data visualization and statistical analyses

Plots were produced using Graphpad Prism. Statistical analyses were performed in R and Graphpad Prism.

## ACKNOWLEDGEMENTS

We thank Susana Godinho for sharing the cell lines. We thank the support of IGC’s Advanced Imaging Unit (AIU), especially Alexandre Lopes, Patrícia Rodrigues and Gabriel Martins, funded by PPBI-POCI-01-0145-FEDER-022122 (Lisboa 2020/FEDER/FCT; Portugal); Flow Cytometry Unit, especially Marta Monteiro, and Advanced Data Analysis Unit, for their support in this work. We thank Raquel Oliveira for insightful and critical discussions, and for her support of C.P. during the final stages of this work. We also thank Elio Sucena and Luca Fava for critical reading of this manuscript and Stephan Peischel for discussion of statistical analyses. We thank the CCR and ED/THEE labs for discussion of this work. This work was supported by the European Research Council (ERC CoG 683528 to M.B.D. and ERC StG 804569 to C.B.) and FCT PhD fellowships awarded to M.L. and C.P. (PD/BD/139217/2018 and PD/BD/128004/2016, respectively).

## AUTHOR CONTRIBUTIONS

M.A.D.L. and C.P. designed and performed all experiments, except for the ones listed below, and discussed all analyses. M.A.D.L. performed centriole number quantifications, quantification of p53 by immunostaining, flow cytometry data acquisition, and all statistical analyses. C.P. performed PLK4 immunostaining and analysis, p53 immunostaining, p53 Western blotting and analysis, and RT-qPCR and analysis. M.S. and C.P. performed the competition assays with the new batch of MCF10A-PLK4, and M.S. performed the rest of the experiment with this new batch, including centriole counting. C.P. and M.A.D.L. supervised M.S.. M.A.D.L., C.P. and M.B.D. critically discussed experimental results. M.A.D.L. and C.P. conceptualised the study. M.A.D.L. and C.P wrote the manuscript. C.B. and M.B.D. edited the manuscript. M.A.D.L. and C.P. contributed equally for this work.

## DECLARATION OF INTERESTS

The authors declare no conflict of interest

## SUPPLEMENTAL INFORMATION

**Figure S1:**
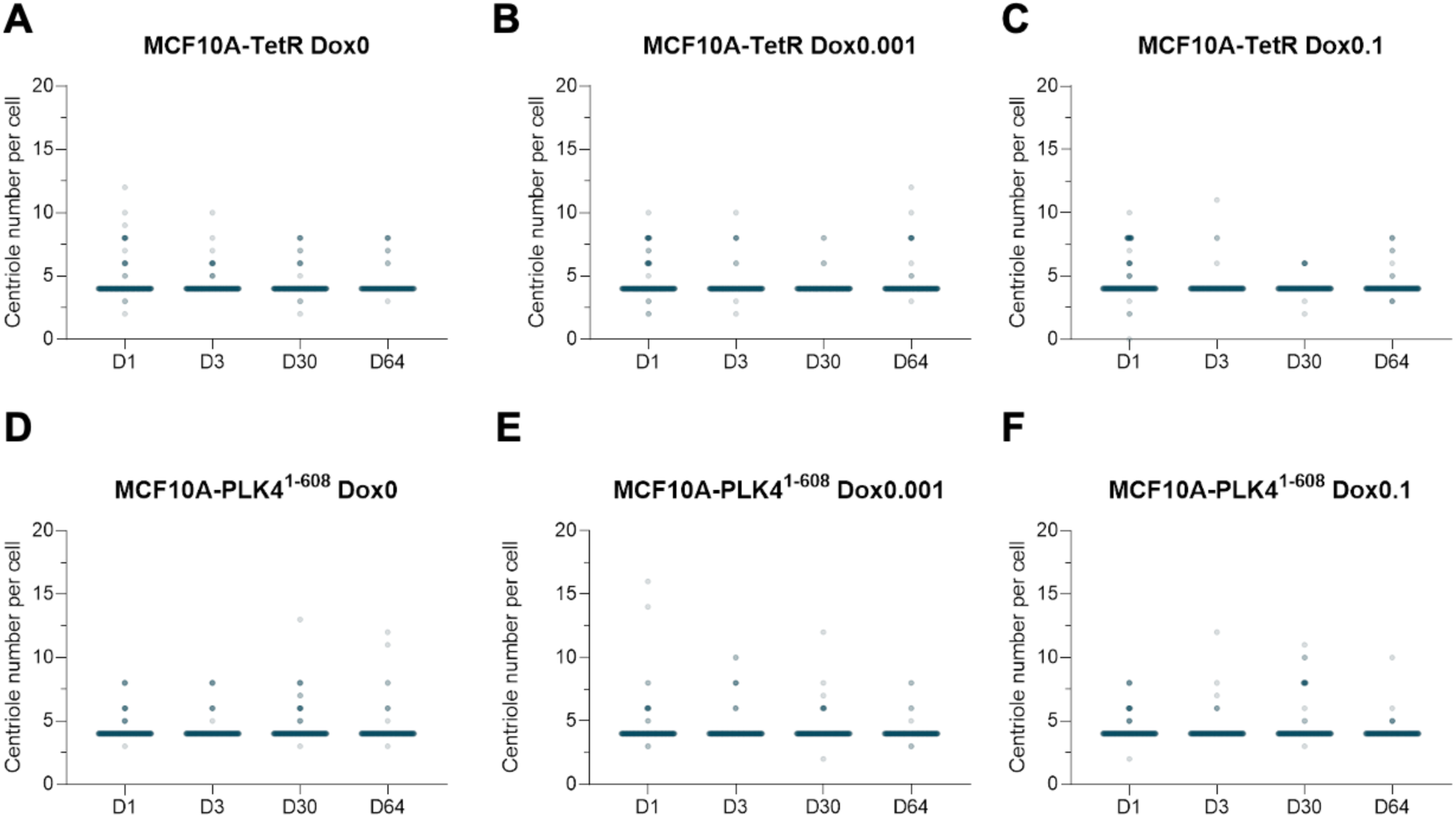
Distributions of centriole numbers per cell in control cell lines. **A-C** - Centriole number distributions in MCF10A-TetR populations grown in Dox0 (A), Dox0.001 (B), and Dox0.1 (C). **D-F** - Centriole number distributions in MCF10A-PLK4^1–608^ populations grown in Dox0 (A), Dox0.001 (B), and Dox0.1 (C). Statistical analysis was performed using linear models for each cell line with Dox concentration and day as independent predictors, plus their interaction, and centriole number as a response. Multiple comparisons with the Dox0-treated population within each day yielded no significant differences at a level ɑ=0.05.

**Figure S2:**
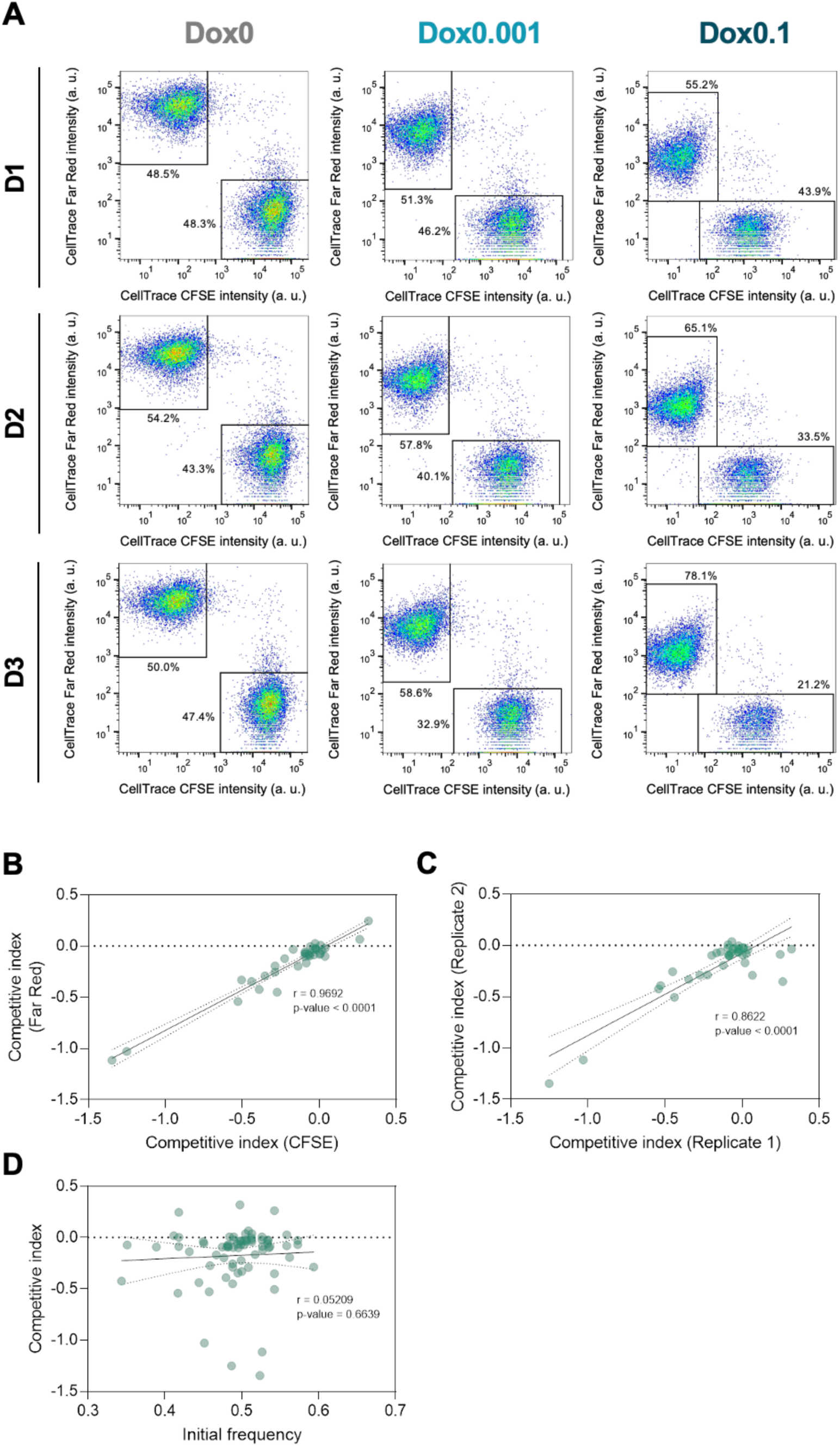
No effect is observed of proliferation dye, replicate and initial frequency on competitive index. **A** - Biparametric plots of MCF10A-Plk4 populations labelled with CellTrace CFSE co-cultured with MCF10A-TetR populations labelled with CellTrace Far Red and treated with Dox0, Dox0.001, and Dox0.1 at days 1, 2, and 3. The percentages of each sub-population is indicated next to the gates. **B** - Correlation between dye combinations in competitive index estimates. CFSE and Far Red refer to the dye which was used to label MCF10A-PLK4 or MCF10A-PLK4^1–608^, in co-culture with MCF10A-TetR. **C** - Correlation between independent experiments in competitive index estimates. **D** - Competitive index estimates as a function of the relative frequency of MCF10A-PLK4 or MCF10A-PLK4^1–608^ at the initial time point of each competition assay (D1, D31, and D64). Data points represent MCF10A-PLK4/MCF10A-PLK4^1–608^ co-cultured with MCF10A-TetR at D1-3, D31-33, and D64-66, grown at Dox0, Dox0.001, or Dox0.1. The black line indicates the best fitting linear regression model. Pearson’s correlation coefficient (*r*) and the *p*-value of the corresponding correlation test indicated in the plots. See Methods for the competitive index expression.

**Figure S3:**
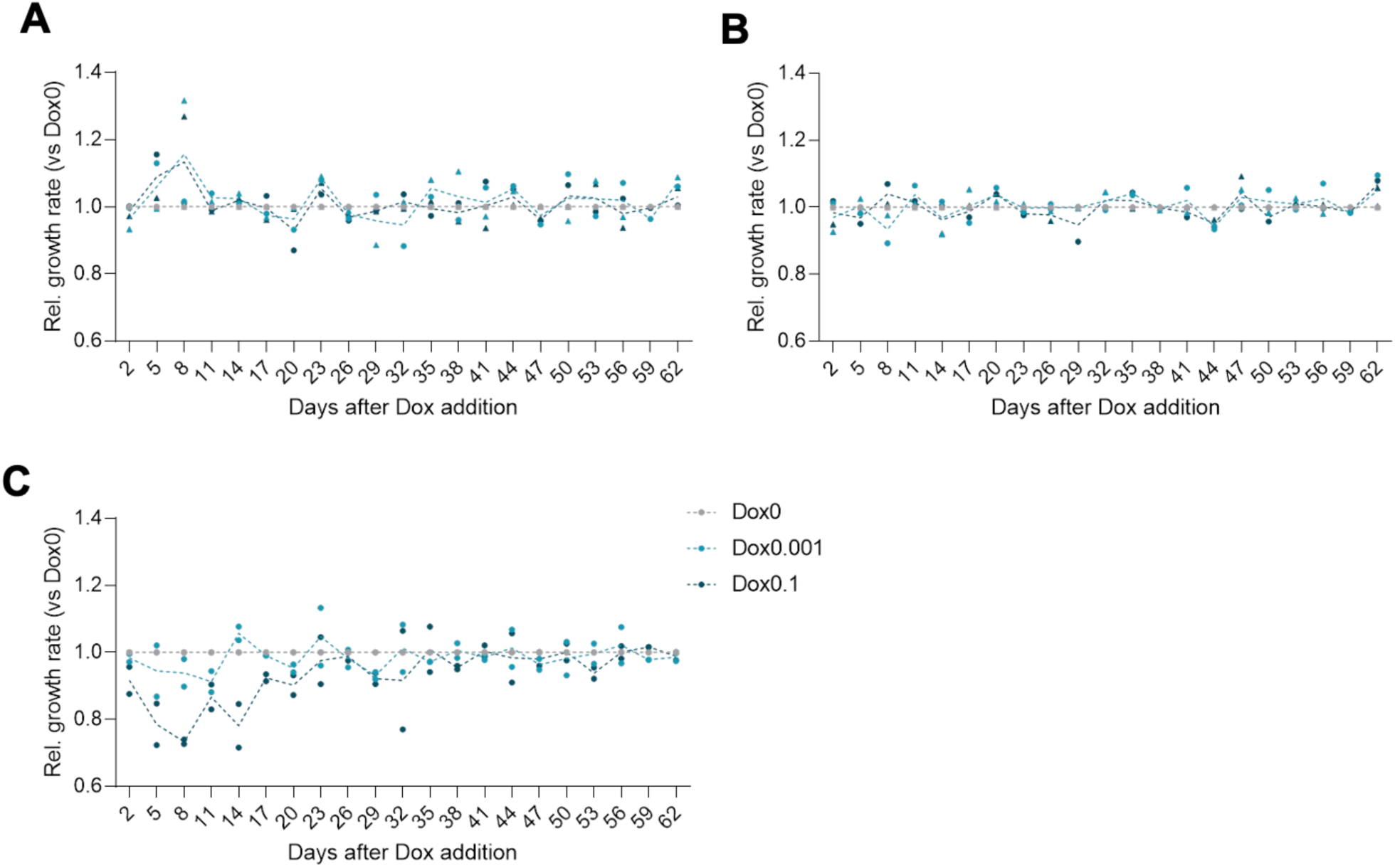
Relative growth rates of each cell line along experimental evolution. **A-C** - Growth rate of MCF10A-TetR (A), MCF10A-PLK4^1–608^(B) and MCF10A-PLK4 (C) treated with Dox0.001 (light blue) or Dox0.1 (dark blue) relative to Dox0 (grey) at the indicated time points along experimental evolution. Data correspond to two independent experiments. See Methods for the relative growth rate expression.

**Figure S4:**
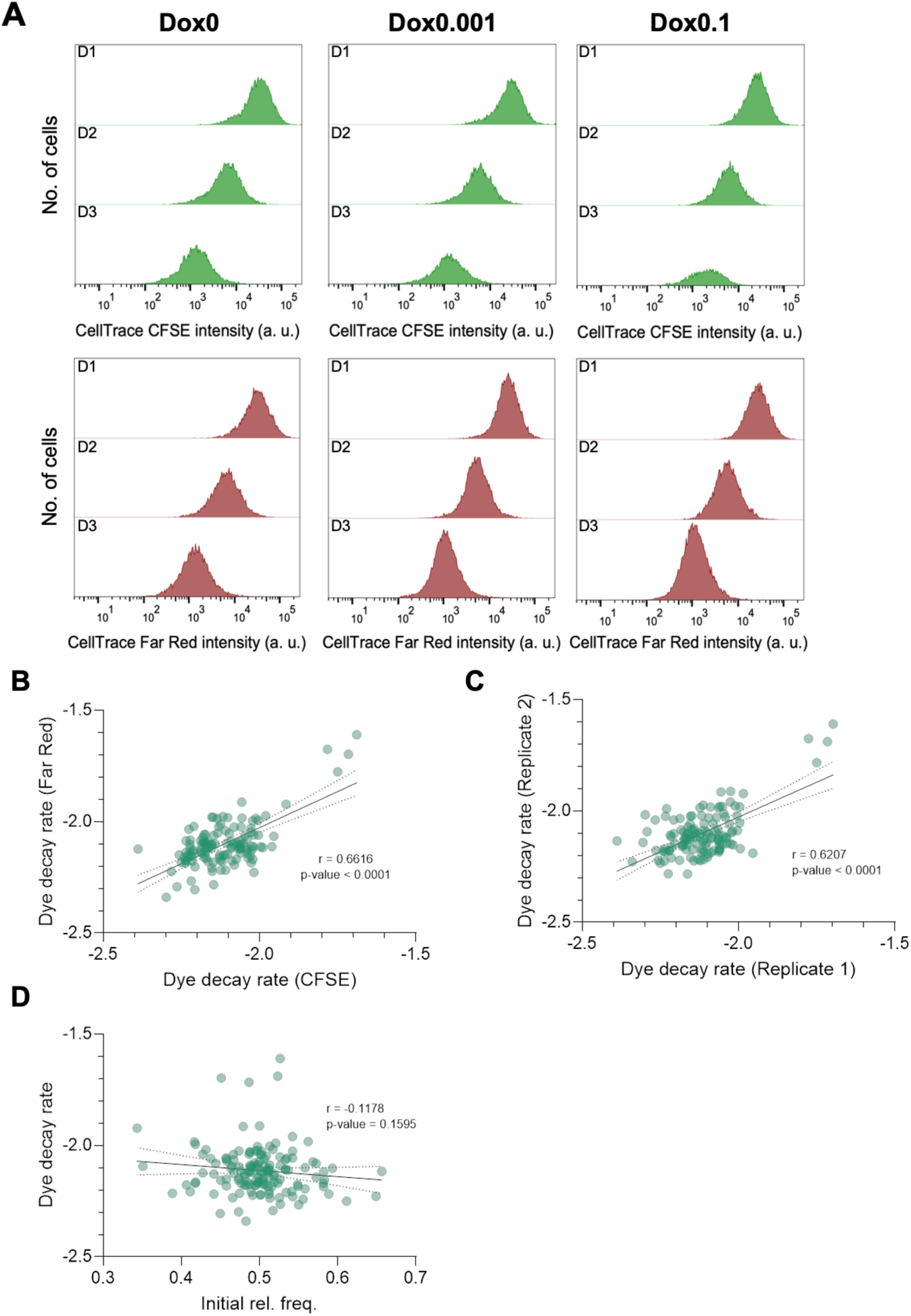
No effect is observed of proliferation dye, replicate and initial frequency on dye decay rates. **A** - Representative histograms of MCF10A-PLK4 labelled with CellTrace CFSE co-cultured MCF10A-TetR labelled with CellTrace Far Red treated with Dox0, Dox0.001, and Dox0.1 at days 1, 2, and 3. **B** – Correlation between dye combinations in dye decay estimates. CFSE and Far Red refer to the dye which was used to label MCF10A-PLK4 or MCF10A-PLK4^1–608^, in co-culture with MCF10A-TetR, or in mono-culture. **C** - Correlation between independent experiments in competitive index estimates. **D** - Competitive index estimates as a function of the relative frequency of MCF10A-PLK4 or MCF10A-PLK4^1–608^ at the initial time point of each competition assay (D1, D31, and D64). Data points represent MCF10A-PLK4/MCF10A-PLK4^1–608^ co-cultured with MCF10A-TetR at D1-3, D31-33, and D64-66, grown at Dox0, Dox0.001, or Dox0.1. The black line indicates the best fitting linear regression model. Pearson’s correlation coefficient (*r*) and the *p*-value of the corresponding correlation test are indicated in the plots. See Methods for the relative dye decay rate expression.

**Figure S5:**
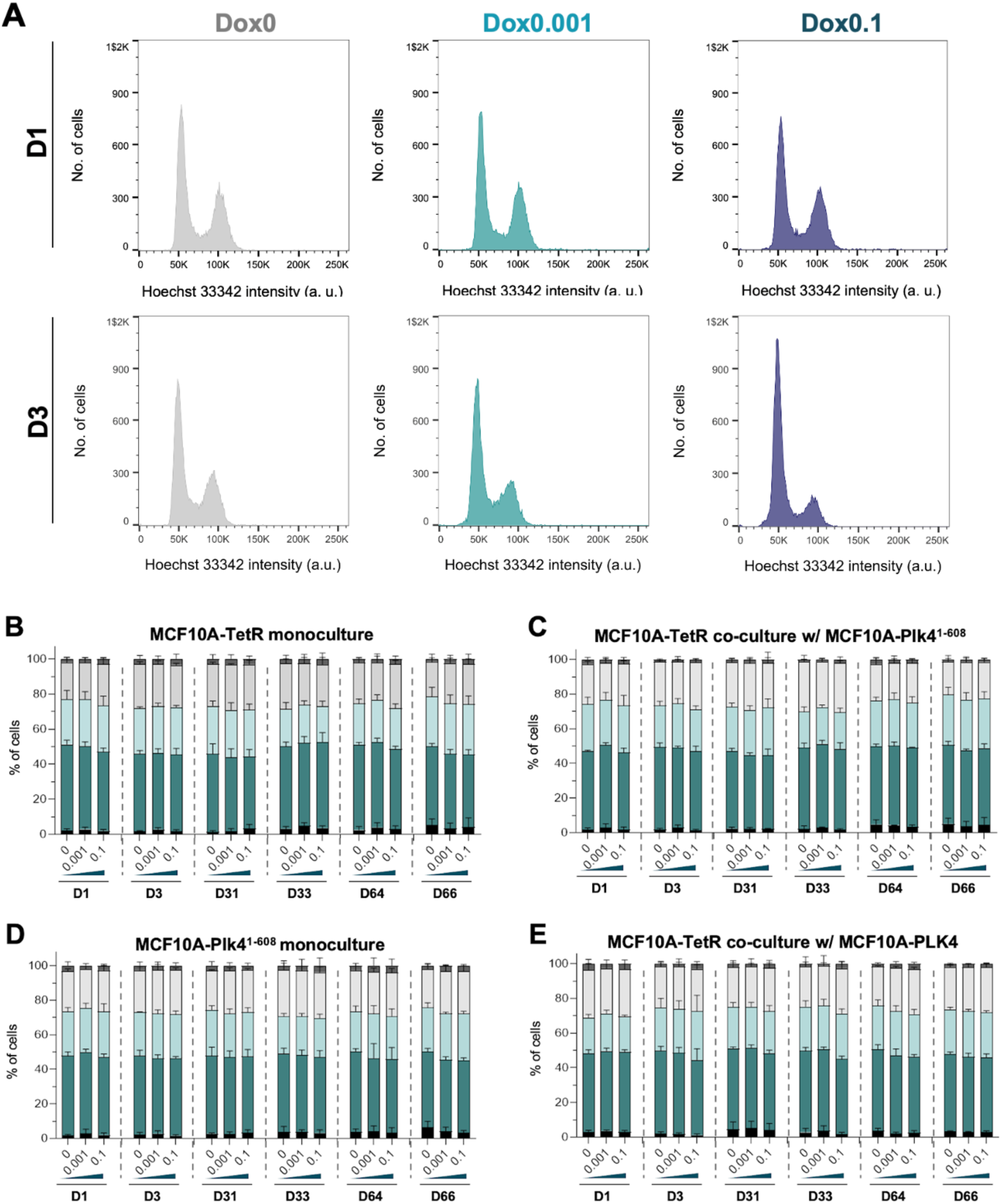
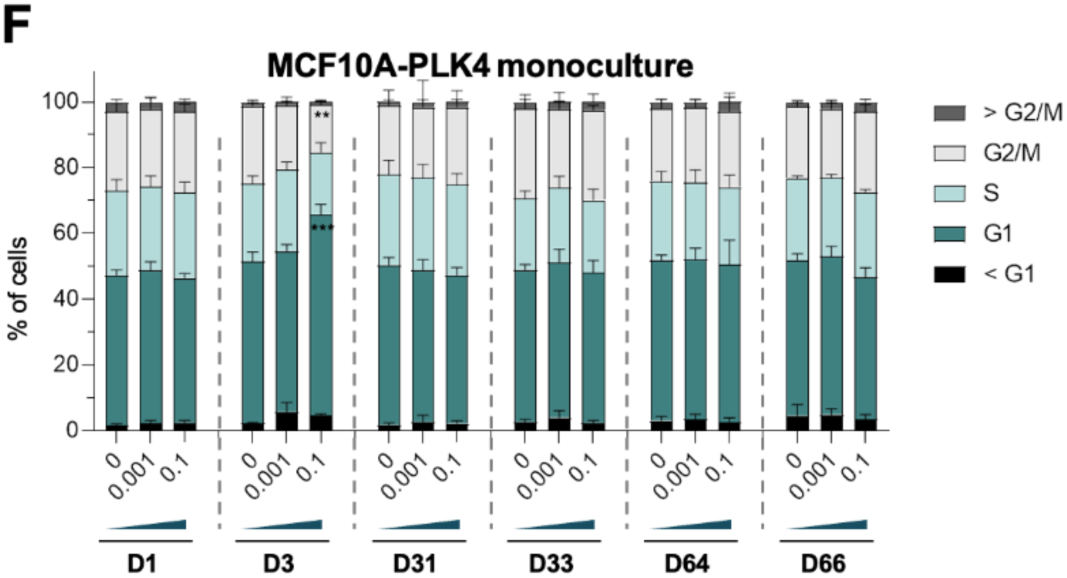
Cell cycle profiles and p53 levels along experimental evolution. **A -** Representative histograms of Hoechst 33342 (DNA) intensity in MCF10A-PLK4 populations treated with Dox0, Dox0.001, and Dox0.1 at days 1 and 3, respectively **B-F -** Relative frequency of cells in G1 (dark green), S (light green), and G2/M (white) for MCF10A-TetR monocultures (B) and co-cultures (C), MCF10A-PLK4^1–608^ monocultures (D) and co-cultures (E) and MCF10A-PLK4 monocultures (F) at the indicated time points. Relative frequencies of co-cultured populations labelled with either dye in the respective cell cycle stage. The error bars represent the mean ± standard deviation. Statistical analysis was performed using generalised linear models taking the percentage of cells in sub-G1, G1, S, G2, or supra-G2 as binomially distributed response variables, with Dox concentration and day as independent predictors, plus their interaction. Significant differences between the indicated cell cycle stages for a given condition and the Dox-0 control at the respective day are represented with *** (*p*-value<0.001).

**Figure S6:**
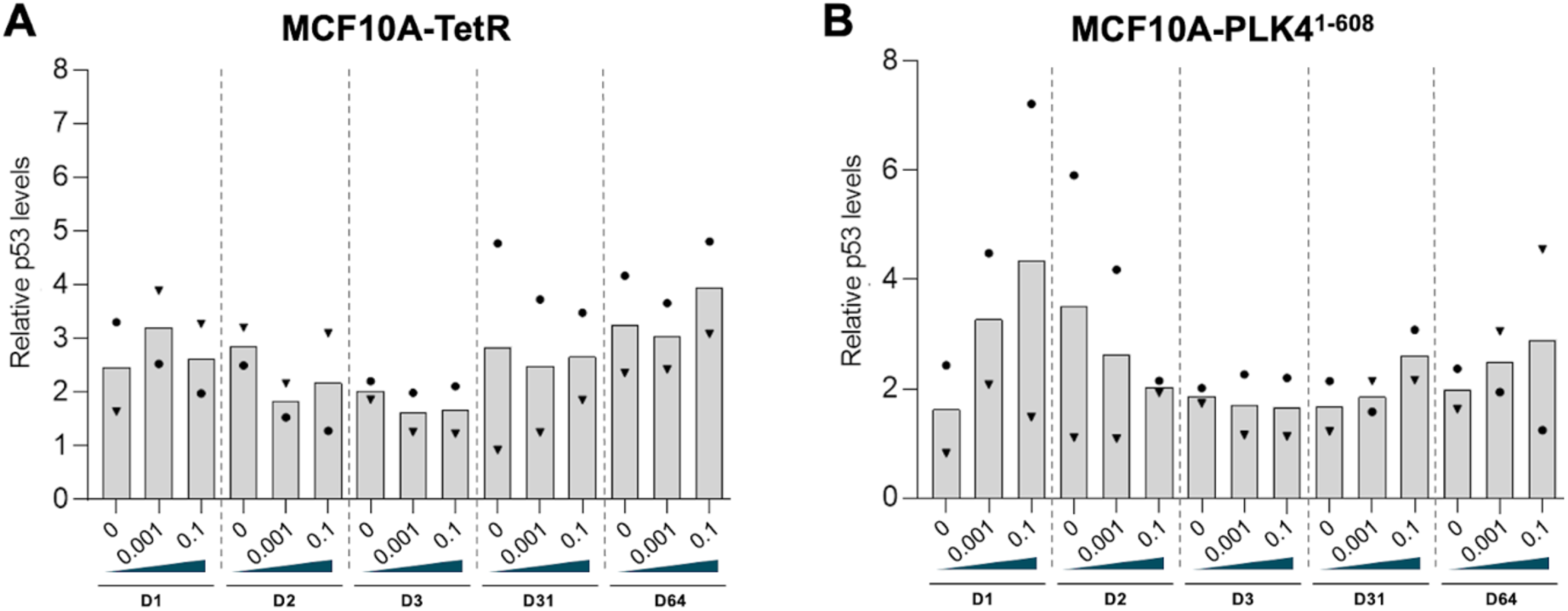
p53 levels along experimental evolution. **A.B -** Quantification of p53 relative to GAPDH by immunoblotting in MCF10A-TetR (A) and MCF10A-PLK4^1–608^ (B) populations treated with the indicated dose of Dox, at the indicated time points. The bars indicate the average between two independent experiments. Statistical analyses were performed using a linear mixed model with Dox concentration and day as independent predictors, plus their interaction, and experiment as a random effect and taking relative p53 levels as a response. Multiple comparisons with the Dox0-treated population within each day yielded no significant differences at a level ɑ=0.05.

**Figure S7:**
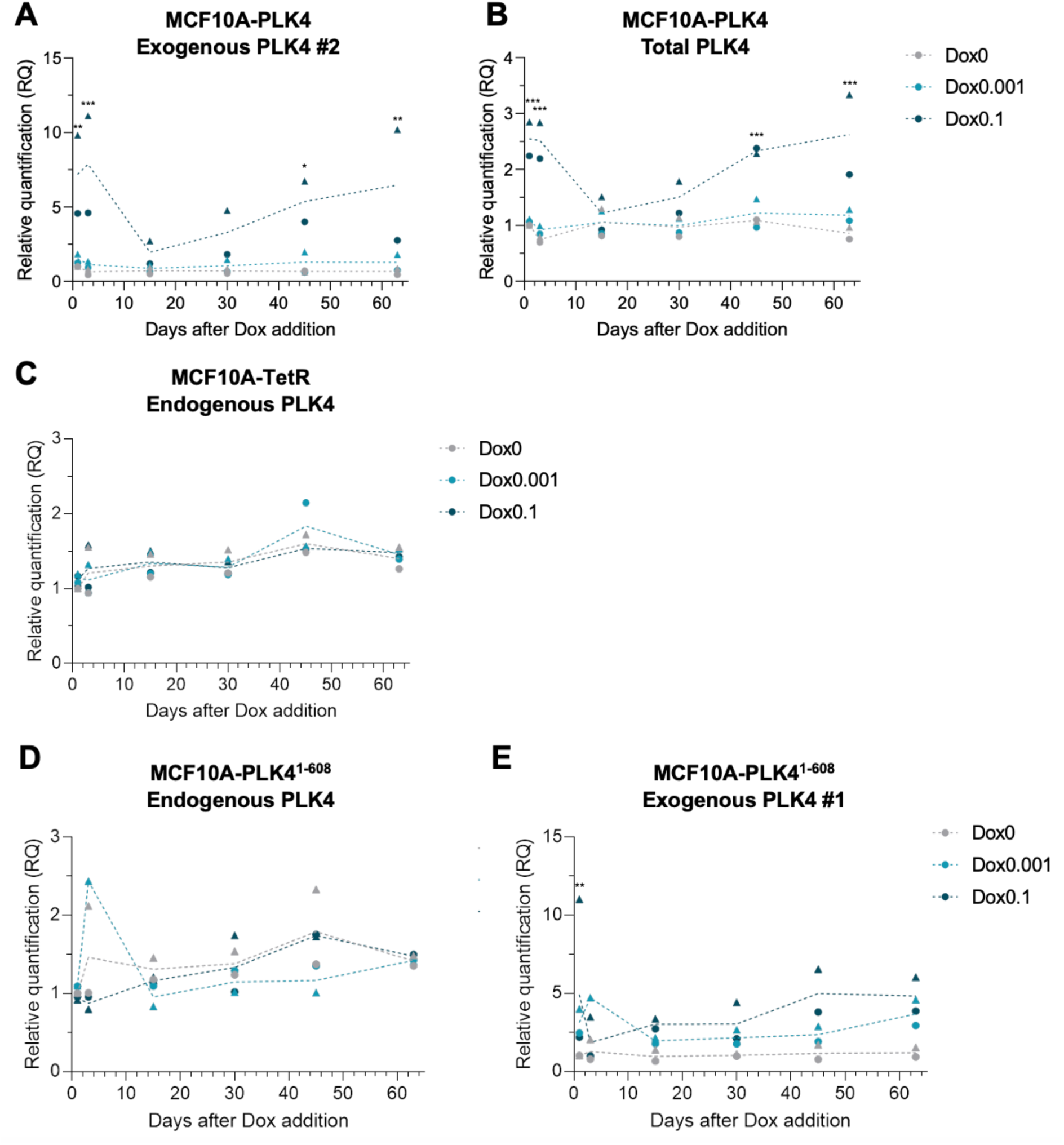
PLK4 expression along experimental evolution. **A-B** - PLK4 mRNA levels in MCF10A-PLK4 analysed by RT-qPCR. The primer sets used are specific for the exogenous PLK4 (A, primer pair #2) or target both the endogenous and exogenous transcripts (B). **C** - Levels of endogenous PLK4 mRNA in MCF10A-TetR at the indicated time points. **D-E** - Levels of endogenous (D) and exogenous (E, primer pair #1) PLK4 mRNA in MCF10A-PLK4^1–608^. The dashed lines correspond to the average RQ across two replicates measured in RNA extracted from MCF10A-PLK4 populations grown with Dox0 (grey), Dox0.001 (blue), and Dox0.1 (dark blue) at the indicated time points and symbols represent different independent experiments. Statistical analyses were performed for each primer set and for each cell line using linear models with Dox concentration and day as independent predictors, plus their interaction, and RQ as a response variable. Significant differences between the indicated conditions and the Dox-0 control at the respective day are represented with *** (*p*-value<0.001).

**Figure S8:**
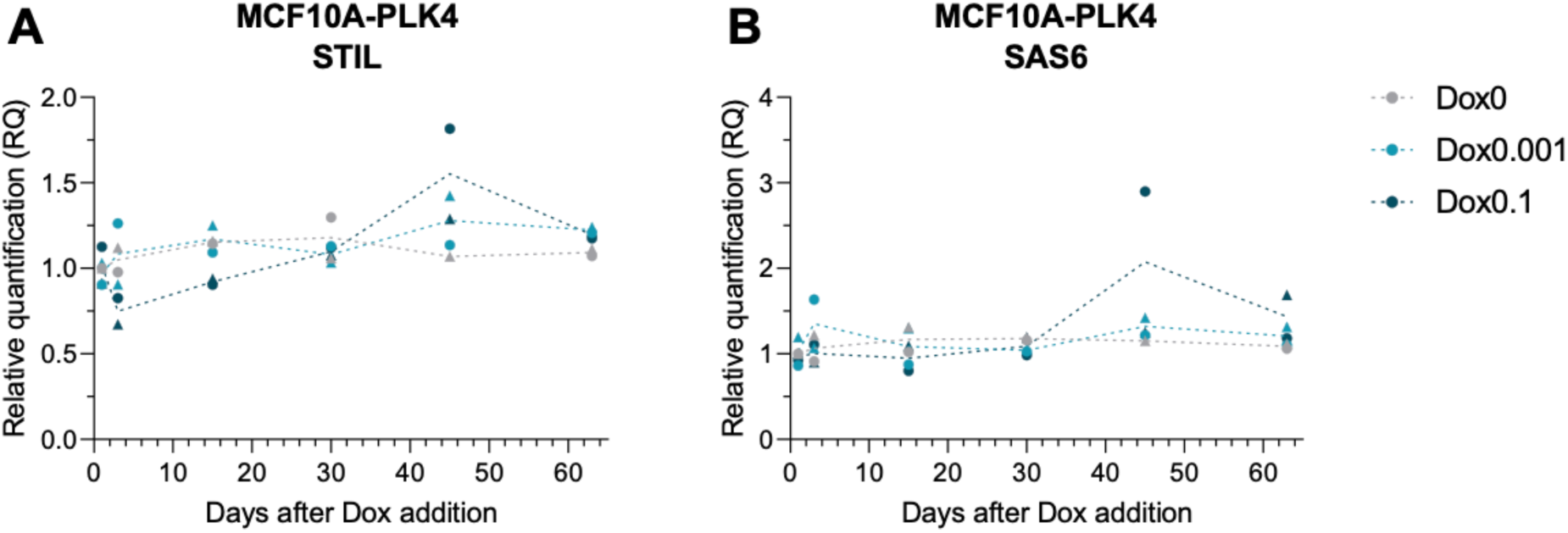
STIL and SAS6 expression along experimental evolution. **A-B** - Levels of STIL (A) and SAS6 (B) mRNA in MCF10A-PLK4 analysed by RT-qPCR. The dashed lines correspond to the average RQ across two independent experiments measured in RNA extracted from MCF10A-PLK4 populations grown with Dox0 (grey), Dox0.001 (blue), and Dox0.1 (dark blue) at the indicated time points. Statistical analyses were performed for each primer set and for each cell line using linear models with Dox concentration and day as independent predictors, plus their interaction, and RQ as a response variable Multiple comparisons with the Dox0-treated population within each day yielded no significant differences at a level ɑ=0.05.

**Table S1:** Predictors of competitive index in CFSE-labelled populations. Results from a linear model taking cell line and doxycycline concentration, and their interaction, as independent variables and competitive index as a response. Data include the competitive index estimated from competition assays performed at days 1-3, 31-33, and 64-66 of two independent evolution experiments when MCF10A-PLK4^1–608^ or MCF10A-PLK4 were labelled with CellTrace CFSE and MCF10A-TetR was labelled with CellTrace Far Red. *β*_0_ - intercept; *β*_MCF10A-PLK4_ - effect of cell line (MCF10A-PLK4 relative to MCF10A-PLK4^1–608^), *β*_Dox0.001_ - effect of 0.001 ug/mL of doxycycline (relative to 0 ug/mL), *β*_Dox0.001_ - effect of 0.1 ug/mL doxycycline (relative to 0 ug/mL), *β*_Dox0.1_ - effect of 0.1 ug/mL of doxycycline (relative to 0 ug/mL), *β*_MCF10A-PLK4*Dox0.001_ - effect of the interaction between cell line MCF10A-PLK4 and doxycycline at 0.001 ug/mL, *β*_MCF10A-PLK4*Dox0.1_ - effect of the interaction between cell line MCF10A-PLK4 and doxycycline at 0.1 ug/mL.

| D1-D3 |  |  |  |  |
| --- | --- | --- | --- | --- |
|  | Estimate | Std. Error | t-value | p-value |
| $\beta_0$ | -0.08007 | 0.031682 | -2.52729 | 0.044844 |
| $\beta_{\text{MCF10A-PLK4}}$ | -0.17627 | 0.044804 | -3.93426 | 0.007675 |
| $\beta_{\text{Dox0.001}}$ | -0.05102 | 0.044804 | -1.13867 | 0.298254 |
| $\beta_{\text{Dox0.1}}$ | -0.01121 | 0.044804 | -0.2501 | 0.810856 |
| $\beta_{\text{MCF10A-PLK4} \times \text{Dox0.001}}$ | -0.16524 | 0.063363 | -2.60775 | 0.040242 |
| $\beta_{\text{MCF10A-PLK4} \times \text{Dox0.1}}$ | -1.03079 | 0.063363 | -16.268 | 3.43E-06 |
| D31-33 |  |  |  |  |
|  | Estimate | Std. Error | t value | p-value |
| $\beta_0$ | -0.00174 | 0.036439 | -0.0478 | 0.963428 |
| $\beta_{\text{MCF10A-PLK4}}$ | -0.09832 | 0.051532 | -1.90799 | 0.105 |
| $\beta_{\text{Dox0.001}}$ | 0.052436 | 0.051532 | 1.017533 | 0.348161 |
| $\beta_{\text{Dox0.1}}$ | -0.05876 | 0.051532 | -1.14018 | 0.29767 |
| $\beta_{\text{MCF10A-PLK4} \times \text{Dox0.001}}$ | -0.23368 | 0.072878 | -3.20652 | 0.018448 |
| $\beta_{\text{MCF10A-PLK4} \times \text{Dox0.1}}$ | -0.30062 | 0.072878 | -4.12505 | 0.006181 |
| D64-66 |  |  |  |  |
|  | Estimate | Std. Error | t value | Pr(>\|t\|) |
| $\beta_0$ | -0.00274 | 0.146459 | -0.01873 | 0.985666 |
| $\beta_{\text{MCF10A-PLK4}}$ | 0.144582 | 0.207124 | 0.698043 | 0.511282 |
| $\beta_{\text{Dox0.001}}$ | -0.03899 | 0.207124 | -0.18826 | 0.856879 |
| $\beta_{\text{Dox0.1}}$ | -0.0044 | 0.207124 | -0.02125 | 0.983734 |
| $\beta_{\text{MCF10A-PLK4} \times \text{Dox0.001}}$ | -0.14786 | 0.292918 | -0.50479 | 0.631707 |

**Table S2:**
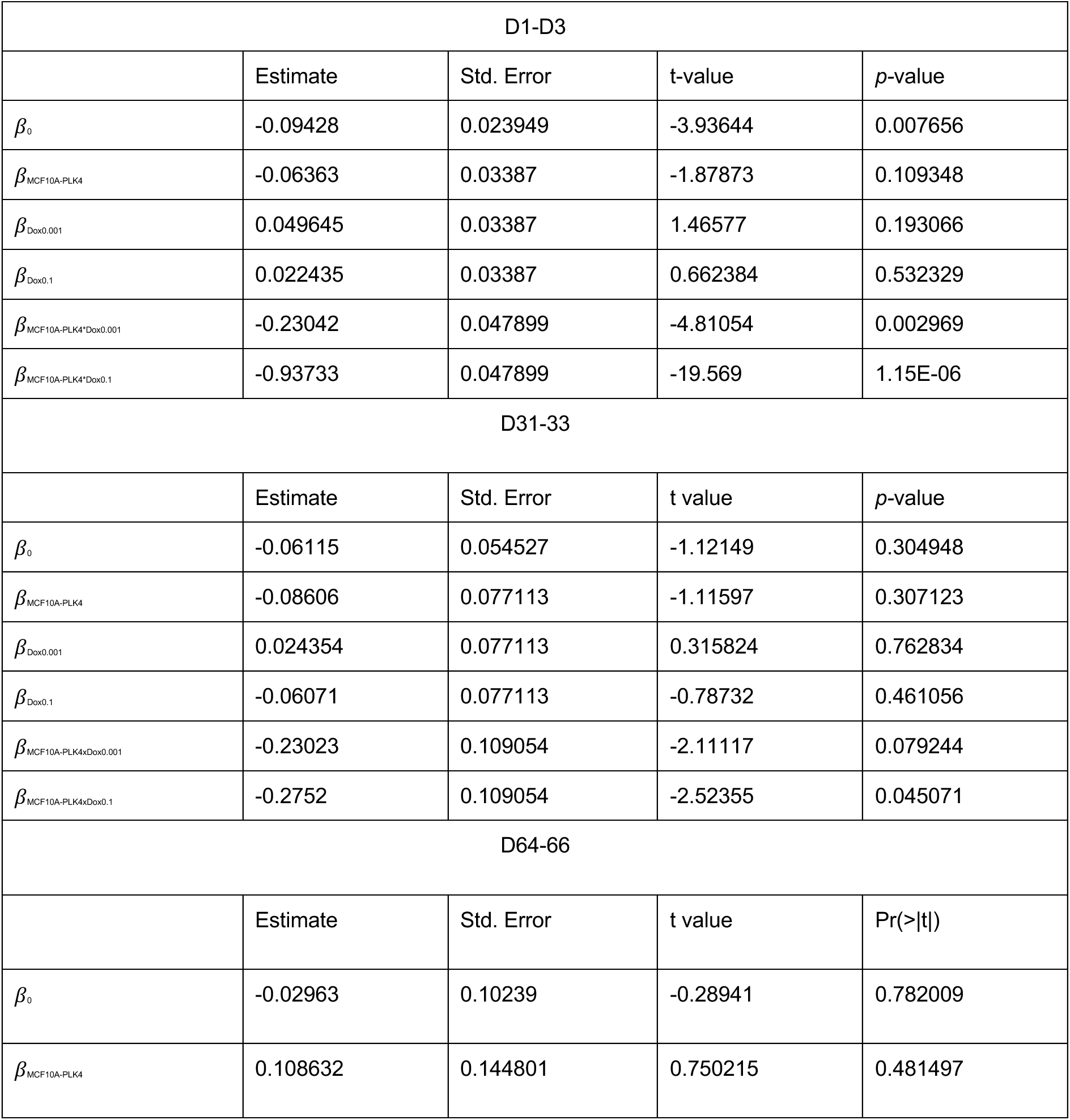
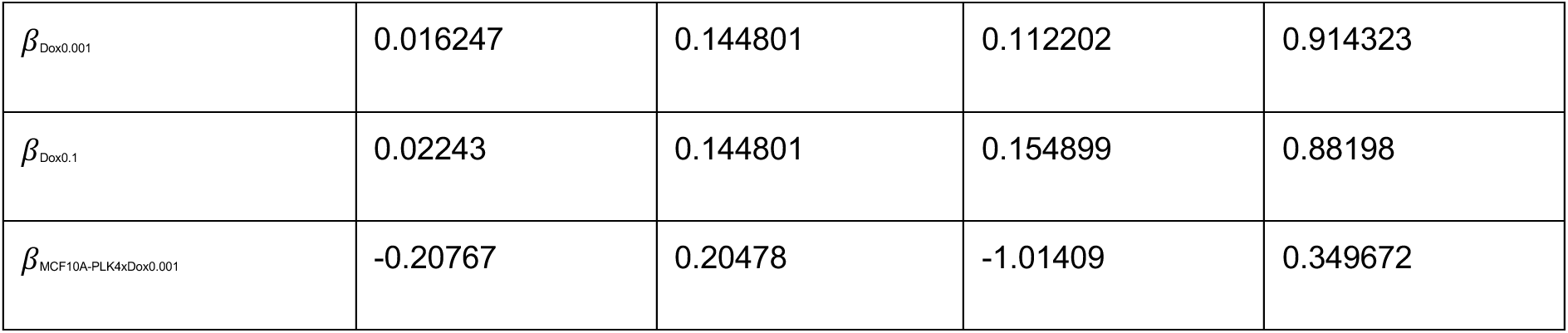
Predictors of competitive index in Far Red-labelled populations. Results from a linear model taking cell line and doxycycline concentration, and their interaction, as independent variables and competitive index as a response. Data include the competitive index estimated from competition assays performed at days 1-3, 31-33, and 64-66 of two independent evolution experiments when MCF10A-PLK4^1–608^ or MCF10A-PLK4 were labelled with CellTrace Far Red and MCF10A-TetR was labelled with CellTrace CFSE. *β*_0_ - intercept; *β*_MCF10A-PLK4_ - effect of cell line (MCF10A-PLK4 relative to MCF10A-PLK4^1–608^), *β*_Dox0.001_ - effect of 0.001 ug/mL of doxycycline (relative to 0 ug/mL), *β*_Dox0.001_ - effect of 0.1 ug/mL doxycycline (relative to 0 ug/mL), *β*_Dox0.1_ - effect of 0.1 ug/mL of doxycycline (relative to 0 ug/mL), *β*_MCF10A-PLK4*Dox0.001_ - effect of the interaction between cell line MCF10A-PLK4 and doxycycline at 0.001 ug/mL, *β*_MCF10A-PLK4*Dox0.1_ - effect of the interaction between cell line MCF10A-PLK4 and doxycycline at 0.1 ug/mL.

| D1-D3 |  |  |  |  |
| --- | --- | --- | --- | --- |
|  | Estimate | Std. Error | t-value | p-value |
| $\beta_0$ | -0.09428 | 0.023949 | -3.93644 | 0.007656 |
| $\beta_{\text{MCF10A-PLK4}}$ | -0.06363 | 0.03387 | -1.87873 | 0.109348 |
| $\beta_{\text{Dox0.001}}$ | 0.049645 | 0.03387 | 1.46577 | 0.193066 |
| $\beta_{\text{Dox0.1}}$ | 0.022435 | 0.03387 | 0.662384 | 0.532329 |
| $\beta_{\text{MCF10A-PLK4} \times \text{Dox0.001}}$ | -0.23042 | 0.047899 | -4.81054 | 0.002969 |
| $\beta_{\text{MCF10A-PLK4} \times \text{Dox0.1}}$ | -0.93733 | 0.047899 | -19.569 | 1.15E-06 |
| D31-33 |  |  |  |  |
|  | Estimate | Std. Error | t value | p-value |
| $\beta_0$ | -0.06115 | 0.054527 | -1.12149 | 0.304948 |
| $\beta_{\text{MCF10A-PLK4}}$ | -0.08606 | 0.077113 | -1.11597 | 0.307123 |
| $\beta_{\text{Dox0.001}}$ | 0.024354 | 0.077113 | 0.315824 | 0.762834 |
| $\beta_{\text{Dox0.1}}$ | -0.06071 | 0.077113 | -0.78732 | 0.461056 |
| $\beta_{\text{MCF10A-PLK4} \times \text{Dox0.001}}$ | -0.23023 | 0.109054 | -2.11117 | 0.079244 |
| $\beta_{\text{MCF10A-PLK4} \times \text{Dox0.1}}$ | -0.2752 | 0.109054 | -2.52355 | 0.045071 |
| D64-66 |  |  |  |  |
|  | Estimate | Std. Error | t value | Pr(>\|t\|) |
| $\beta_0$ | -0.02963 | 0.10239 | -0.28941 | 0.782009 |
| $\beta_{\text{MCF10A-PLK4}}$ | 0.108632 | 0.144801 | 0.750215 | 0.481497 |
| $\beta_{Dox0.001}$ | 0.016247 | 0.144801 | 0.112202 | 0.914323 |
| $\beta_{Dox0.1}$ | 0.02243 | 0.144801 | 0.154899 | 0.88198 |
| $\beta_{MCF10A-PLK4 \times Dox0.001}$ | -0.20767 | 0.20478 | -1.01409 | 0.349672 |

**Table S3:** Predictors of relative dye dilution rate in co-cultured CFSE-labelled populations. Results from a linear model taking cell line and doxycycline concentration, and their interaction, as independent variables and relative dye dilution rate as a response. Data include the competitive index estimated from competition assays performed at days 1-3, 31-33, and 64-66 of two independent evolution experiments when MCF10A-PLK4^1–608^ or MCF10A-PLK4 were labelled with CellTrace CFSE and MCF10A-TetR was labelled with CellTrace Far Red. *β*_0_ - intercept; *β*_MCF10A-PLK4_ - effect of cell line (MCF10A-PLK4 relative to MCF10A-PLK4^1–608^), *β*_Dox0.001_ - effect of 0.001 ug/mL of doxycycline (relative to 0 ug/mL), *β*_Dox0.001_ - effect of 0.1 ug/mL doxycycline (relative to 0 ug/mL), *β*_Dox0.1_ - effect of 0.1 ug/mL of doxycycline (relative to 0 ug/mL), *β*_MCF10A-PLK4*Dox0.001_ - effect of the interaction between cell line MCF10A-PLK4 and doxycycline at 0.001 ug/mL, *β*_MCF10A-PLK4*Dox0.1_ - effect of the interaction between cell line MCF10A-PLK4 and doxycycline at 0.1 ug/mL.

| D1-D3 |  |  |  |  |
| --- | --- | --- | --- | --- |
|  | Estimate | Std. Error | t-value | p-value |
| $\beta_0$ | 1.034523 | 0.012241 | 84.51014 | 1.85E-10 |
| $\beta_{MCF10A-PLK4}$ | -0.02174 | 0.017312 | -1.25563 | 0.255935 |
| $\beta_{Dox0.001}$ | 0.002239 | 0.017312 | 0.129332 | 0.901322 |
| $\beta_{Dox0.1}$ | -0.00239 | 0.017312 | -0.13808 | 0.894698 |
| $\beta_{MCF10A-PLK4 \times Dox0.001}$ | -0.0372 | 0.024483 | -1.51948 | 0.179448 |
| $\beta_{MCF10A-PLK4 \times Dox0.1}$ | -0.21281 | 0.024483 | -8.69221 | 0.000128 |
| D31-33 |  |  |  |  |
|  | Estimate | Std. Error | t value | p-value |
| $\beta_0$ | 0.998159 | 0.006573 | 151.8668 | 5.50E-12 |
| $\beta_{MCF10A-PLK4}$ | -0.01745 | 0.009295 | -1.87737 | 0.109554 |
| $\beta_{Dox0.001}$ | 0.010539 | 0.009295 | 1.133832 | 0.300126 |
| $\beta_{Dox0.1}$ | -0.01967 | 0.009295 | -2.11588 | 0.078731 |
| $\beta_{MCF10A-PLK4 \times Dox0.001}$ | -0.04343 | 0.013145 | -3.30389 | 0.016329 |
| $\beta_{\text{MCF10A-PLK4} \times \text{Dox0.1}}$ | -0.04813 | 0.013145 | -3.66109 | 0.010566 |
| D64-66 |  |  |  |  |
|  | Estimate | Std. Error | t value | Pr(>\|t\|) |
| $\beta_0$ | 0.960969 | 0.021168 | 45.39803 | 7.65E-09 |
| $\beta_{\text{MCF10A-PLK4}}$ | -0.02529 | 0.029936 | -0.84472 | 0.430657 |
| $\beta_{\text{Dox0.001}}$ | 0.011394 | 0.029936 | 0.380602 | 0.716609 |
| $\beta_{\text{Dox0.1}}$ | 0.018177 | 0.029936 | 0.607212 | 0.565965 |
| $\beta_{\text{MCF10A-PLK4} \times \text{Dox0.001}}$ | -0.02888 | 0.042335 | -0.68206 | 0.520649 |

**Table S4:** Predictors of relative dye dilution rate in co-cultured Far Red-labelled populations. Results from a linear model taking cell line and doxycycline concentration, and their interaction, as independent variables and relative dye dilution rate as a response. Data include the competitive index estimated from competition assays performed at days 1-3, 31-33, and 64-66 of two independent evolution experiments when MCF10A-PLK4^1–608^ or MCF10A-PLK4 were labelled with CellTrace Far Red and MCF10A-TetR was labelled with CellTrace CFSE. *β*_0_ - intercept; *β*_MCF10A-PLK4_ - effect of cell line (MCF10A-PLK4 relative to MCF10A-PLK4^1–608^), *β*_Dox0.001_ - effect of 0.001 ug/mL of doxycycline (relative to 0 ug/mL), *β*_Dox0.001_ - effect of 0.1 ug/mL doxycycline (relative to 0 ug/mL), *β*_Dox0.1_ - effect of 0.1 ug/mL of doxycycline (relative to 0 ug/mL), *β*_MCF10A-PLK4*Dox0.001_ - effect of the interaction between cell line MCF10A-PLK4 and doxycycline at 0.001 ug/mL, *β*_MCF10A-PLK4*Dox0.1_ - effect of the interaction between cell line MCF10A-PLK4 and doxycycline at 0.1 ug/mL.

|  |  |  |  |  |
| --- | --- | --- | --- | --- |
| D1-D3 |  |  |  |  |
|  | Estimate | Std. Error | t-value | p-value |
| $\beta_0$ | 0.946771 | 0.009657 | 98.04326 | 7.59E-11 |
| $\beta_{\text{MCF10A-PLK4}}$ | -0.02093 | 0.013657 | -1.53287 | 0.176197 |
| $\beta_{\text{Dox0.001}}$ | 0.008626 | 0.013657 | 0.631618 | 0.550927 |
| $\beta_{\text{Dox0.1}}$ | 0.016351 | 0.013657 | 1.197298 | 0.276343 |
| $\beta_{\text{MCF10A-PLK4} \times \text{Dox0.001}}$ | -0.04444 | 0.019313 | -2.30078 | 0.061036 |
| $\beta_{\text{MCF10A-PLK4} \times \text{Dox0.1}}$ | -0.19201 | 0.019313 | -9.94196 | 5.99E-05 |
| D31-33 |  |  |  |  |
|  | Estimate | Std. Error | t value | p-value |
| $\beta_0$ | 1.006421 | 0.016872 | 59.6513 | 1.49E-09 |
| $\beta_{\text{MCF10A-PLK4}}$ | -0.0203 | 0.02386 | -0.85082 | 0.427512 |
| $\beta_{\text{Dox0.001}}$ | -0.00107 | 0.02386 | -0.04475 | 0.965762 |
| $\beta_{\text{Dox0.1}}$ | 0.004794 | 0.02386 | 0.200918 | 0.8474 |
| $\beta_{\text{MCF10A-PLK4xDox0.001}}$ | -0.03262 | 0.033743 | -0.96666 | 0.371046 |
| $\beta_{\text{MCF10A-PLK4xDox0.1}}$ | -0.04681 | 0.033743 | -1.38727 | 0.214692 |
| D64-66 |  |  |  |  |
|  | Estimate | Std. Error | t value | Pr(>\|t\|) |
| $\beta_0$ | 1.018748 | 0.02016 | 50.53233 | 4.03E-09 |
| $\beta_{\text{MCF10A-PLK4}}$ | 0.00492 | 0.028511 | 0.172555 | 0.868675 |
| $\beta_{\text{Dox0.001}}$ | 0.014508 | 0.028511 | 0.508856 | 0.629018 |
| $\beta_{\text{Dox0.1}}$ | 0.013561 | 0.028511 | 0.475656 | 0.651143 |
| $\beta_{\text{MCF10A-PLK4xDox0.001}}$ | -0.05015 | 0.040321 | -1.24389 | 0.259932 |

## Notes

### Competing Interest Statement

The authors have declared no competing interest.

### Summary of Updates

Updated with new results (last figure) and an extra author who contributed to the new experiments.

